# Evolution of selfish multicellularity collective organisation of individual spatio-temporal regulatory strategies

**DOI:** 10.1101/2022.04.08.487695

**Authors:** Renske M.A. Vroomans, Enrico Sandro Colizzi

**Affiliations:** Origins Center, Netherlands; Informatics Institute, University of Amsterdam; Origins Center, Netherlands; Mathematical Institute, Leiden University; Sainsbury Laboratory, University of Cambridge

**Keywords:** Evolution of multicellularity, evolution of regulation, computational modelling

## Abstract

**Background:** The unicellular ancestors of modern-day multicellular organisms were remarkably complex. They had an extensive set of regulatory and signalling genes, an intricate life cycle and could change their behaviour in response to environmental changes. At the transition to multicellularity, some of these behaviours were co-opted to organise the development of the nascent multicellular organism. Here, we focus on the transition to multicellularity before the evolution of stable cell differentiation, to reveal how the emergence of clusters affects the evolution of cell behaviour.

**Results:** We construct a computational model of a population of cells that can evolve the regulation of their behavioural state – either division or migration – in a unicellular or multicellular context. They compete for reproduction and for resources to survive in a seasonally changing environment. We find that the evolution of multicellularity strongly determines the co-evolution of cell behaviour, by altering the competition dynamics between cells. When adhesion cannot evolve, cells compete for survival by rapidly migrating towards resources before dividing. When adhesion evolves, emergent collective migration alleviates the pressure on individual cells to reach resources. This allows cells to selfishly maximise replication. Migrating adhesive clusters display striking patterns of spatio-temporal cell state changes that visually resemble animal development.

**Conclusions:** Our model demonstrates how emergent selection pressures at the onset of multicellularity can drive the evolution of cellular behaviour to give rise to developmental patterns.

## Background

The evolution of multicellularity is a major transition in individuality, which occurred multiple times across the tree of life [1–3]. These transitions were likely driven by an initial increase in cell-cell adhesion [4] - as also shown by *in vitro* evolution experiments [5, 6], leading to cluster formation by aggregation or by inhibition of cell separation [3, 7–9]. The unicellular ancestors of these nascent multicellular organisms exhibited complex behaviour and were capable of switching to different phenotypes in response to changes in the environment [10]: a form of reversible differentiation. When adhesion evolved, the newly multicellular aggregates consisted of complex cells that could exploit their pre-existing ability to differentiate in this new biotic context. Eventually the genetic toolkit organising differentiation gave rise to the developmental program of complex multicellular organisms, with irreversible cell differentiation organised through spatio-temporal pattern formation [3, 11, 12].

The evolution of cell differentiation at the onset of multicellularity has been studied in computational models from the perspective of division of labour (usually between soma and germ line), where trade-offs between costs and benefits of functional specialisation and multicellularity determine the size of clusters [13–16]. These models assume the existence of multiple stable cell states, and find the selective conditions under which both states arise in a group of cells. Spatial differentiation patterns and cell-cell communication can also naturally arise as a means to stabilise intracellular gene regulation [17], or as a by-product of selection for different cell-types [18]. Alternatively, multicellular life-cycles can arise spontaneously because of conflicts between social groups of cells [19], or because of the interactions between cells’ internal regulation and a changing environment [20]. Group dynamics emerging in spatially structured models can drive the evolution of differentiation, resulting in proto-developmental dynamics [21] and genome structuring [22]. However, it is unclear how the temporal regulatory program of the unicellular ancestors gave rise to the developmental program of multicellular organisms.

Here, we construct a computational model to investigate the first steps of this evolutionary transition – from the ecology of unicellular populations to that of nascent multicellular organisms, prior to the evolution of stable cell differentiation. Specifically, we set out to show how the evolution of adhering cell clusters impacts the evolution of cell behaviour regulation. We then show how this results in the emergence of coordinated development-like dynamics (i.e. without imposing an external selection pressure). Finally, we characterise the evolutionary dynamics of both the unicellular and the multicellular state.

In the model, an evolving population of spatially embedded cells have to find resources to survive. Cells have evolvable adhesion proteins, allowing for a spectrum of adhesion strength, and an evolvable regulatory network which determines their behaviour in response to a seasonally changing environment. Through their regulatory network, cells decide when to divide, and when to migrate towards the resources, allowing for various survival strategies that are characterised by one or multiple phenotypic switches between these two states.

We find that the evolved strategy depends on whether adhesion can co-evolve, especially when the environment imposes a high selection pressure. When adhesion cannot evolve, competition between cells is dominated by reaching the resources first – they have to navigate the abiotic environment to find the resources by themselves. When adhesion can evolve, cells can perform collective migration. This lowers the pressure to reach resources quickly, because cells carry each other to the peak of the gradient [23]. Competition between cells then becomes driven by ensuring more division. This maximization of individual’s reproductive success in the multicellular group shows that selection on cell behaviour becomes dominated by the newly established biotic environment. Thus, even within the simple context of our model, we observe a complex evolutionary transient that involves intricate feedback between cooperation (in the form of collective migration) and competition for survival and reproduction. The interactions between cells within clusters, and between cells and the environment, lead to an emergent coordination of migration and division which strongly resembles developmental processes.

## Results

### Model setup

We model an evolving population of cells that have to locate resources necessary for survival in a periodically changing environment, throughout multiple seasons. We implement a 2D hybrid Cellular Potts Model (CPM) [24–26] for the cellular dynamics, using a square lattice for the cell population and a lattice of the same dimensions for a chemoattractant signal (Fig. 1A). CPM has been extensively used to model many aspects of embryonic development [27–29], since it endows cells with an explicit size and shape, allowing for both subcellular resolution and deformation, as well as cell level properties such as adhesion and migration. We have previously shown that CPM is very suitable to modelling uni-cellular eco-evolutionary dynamics at the transition to multicellularity [23].

**Figure 1:**
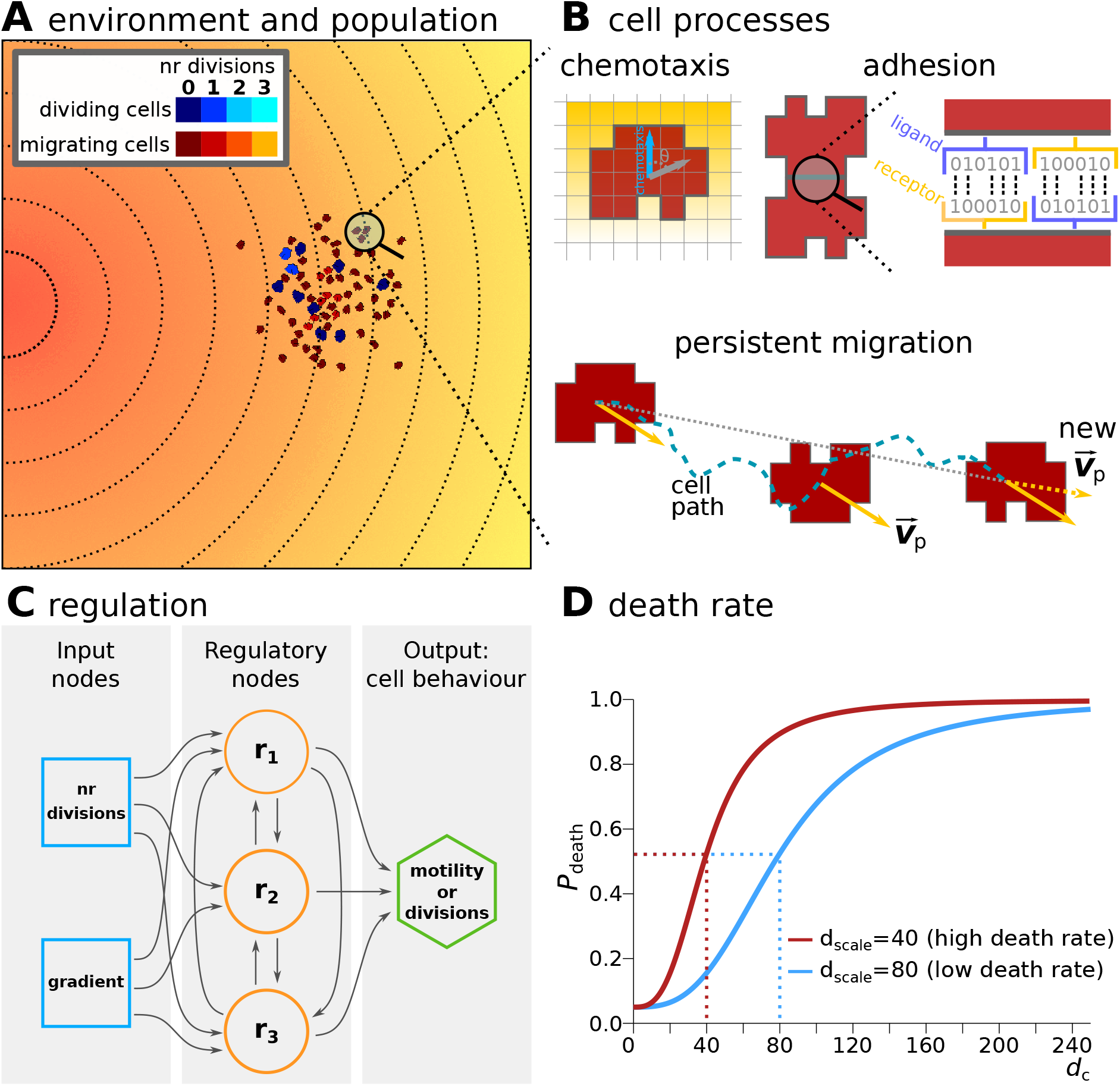
Model description. **a)** The environment in which cells have to survive contains a chemoattractant gradient (lines and colour indicate equal amounts of chemoattractant). **b)** Cells can sense the chemoattractant in the lattice sites that correspond to their own location, and move preferentially in the direction of perceived higher concentration (the blue arrow). Adhesion between two cells is mediated by receptors and ligands (represented by a bitstring, see Methods). The receptor of one cell is matched to the ligand of the other cell and vice versa. The more complementary the receptors and ligands are, the lower the J values and the stronger the adhesion between the cells. Persistent migration is implemented by endowing each cell with a preferred direction of motion 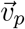. Every *τ_p_* MCS, this direction is updated with a cell’s actual direction of motion in that period. **c)** Cells have a simple evolvable regulatory network to determine whether to migrate or divide at any given time (with max 3 divisions). The network receives as input the number of divisions the cell has already done, and the concentration of the gradient. The activation threshold (*ρ_i_*) of each node *i* and strength of interaction (*w_j_*) of node *j* on node *i* can evolve to have a different effect on the state *S_i_* of the node. **d)** Probability for a cell to die at the end of a season, as a function of its distance to the peak *d_c_* (in lattice sites). *d*_scale_ determines the distance at which the probability is half-maximal.

In the current model, cells can adhere to each other based on the ligands and receptors that they express on their membrane (Fig. 1B). The greater the complementarity between the ligands and receptors of two cells that are in contact, the stronger their adhesion. This is translated to the cell-cell and cell-medium adhesion energy in the CPM (respectively *J*_c,c_ and *J*_c,m_, see Methods). The adhesion strength resulting from *J*_c,c_ and *J*_c,m_ is quantified by the surface tension 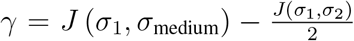. Cells adhere to each other when *γ* > 0 (i.e. with lower contact energy), while they preferentially interact with the medium - and thus do not adhere - when *γ* <0 θ (*γ* = 0 corresponds to neutral adhesion). In some simulations, the receptors and ligands can evolve, thereby changing the strength of adhesion between cells. In simulations where adhesion cannot evolve, we set *J*_c,c_ and *J*_c,m_ such that the surface tension *γ* between cells and the medium is negative, resulting in no adhesion.

Cells can be in one of two states: migratory or dividing, which are mutually exclusive phe-notypes in our model (due to conflicting use of the cytoskeleton). When the *in silico* cells are migratory, they perform a persistent random walk and chemotaxis towards higher local concentrations of the chemoattractant signal (Fig. 1B). The chemoattractant is deposited as a gradient with a peak on one side of the lattice; this gradient is steep enough that cells are able to migrate to the peak individually. The location of the peak changes to a random side of the lattice (up, down, left or right) at the start of each new season. Dividing cells are stationary and do not react to the chemoattractant. When cells have been in the “dividing” state continuously for 10000 CPM update steps (Monte Carlo Steps, MCS – see Methods), they slowly start increasing their size over another 20000 MCS. Cells divide when they have grown to twice their original size. Each cell can divide a maximum of three times during each season, implicitly assuming that resources acquired by a cell are finite, and a cell’s available energy is capped.

The state of a cell is determined by an evolvable gene regulatory network (GRN), modeled as a Boolean network with a fixed architecture, loosely based on [30] (Fig. 1C). The network receives two inputs: the average local concentration of the chemoattractant at the cell’s location, and the number of times the cell has already divided. It has three regulatory genes that process the input and determine the state of the output gene. The state of the output gene in turn determines cell state: migratory or dividing (see Methods). When a cell has performed the maximum number of divisions in a season, it cannot divide anymore even when the state of its output gene dictates a dividing state. Instead, the cell is in a quiescent state in which it is neither dividing nor migrating. When a cell divides, the daughter cell inherits the receptors and ligands for adhesion and the GRN architecture; the state of all genes in the networks of both cells is reset to 0. In one of the daughter cells, mutations can happen in the strength of the gene regulatory interactions in the GRN and in the activation threshold of each gene.

Each season lasts a fixed number of MCS, so that cells have a fixed amount of time to divide and migrate. Different simulations can have seasons of different lengths. Cells that are closer to the peak of the gradient at the end of the season have a higher probability to survive into the next season than cells that are further away (Fig. 1D). At the start of each new season, the division counter in each cell, and the states of the genes in their GRN, are reset to 0.

### Short seasons and high death rates select for polarised regulatory strategies

We assessed the effect of season duration, seasonal death rate and evolution of adhesion on the evolution of cells’ regulatory strategy. We ran 15 simulations for each combination of season duration (short, intermediate or long), seasonal death rate (low or high), and the possibility of evolving adhesion, as summarised in Table 1. Each simulation was initialised with a starting population of cells possessing random GRNs, which could evolve over subsequent seasons. For milder conditions (lower death rates and longer seasons), populations in all 15 simulations could evolve viable strategies, in which cells switch at least once between migration and dividing (Fig. 2A). This led to a large population capable of reaching the peak of the gradient before the end of the season. In the simulations with intermediate season duration, higher death rates caused some populations to go extinct because cells were unable to evolve such switching. Under the harshest conditions (high death rates and very short seasons), extinction occurred in 4 to 7 simulation replicas (table 1).

**Figure 2:**
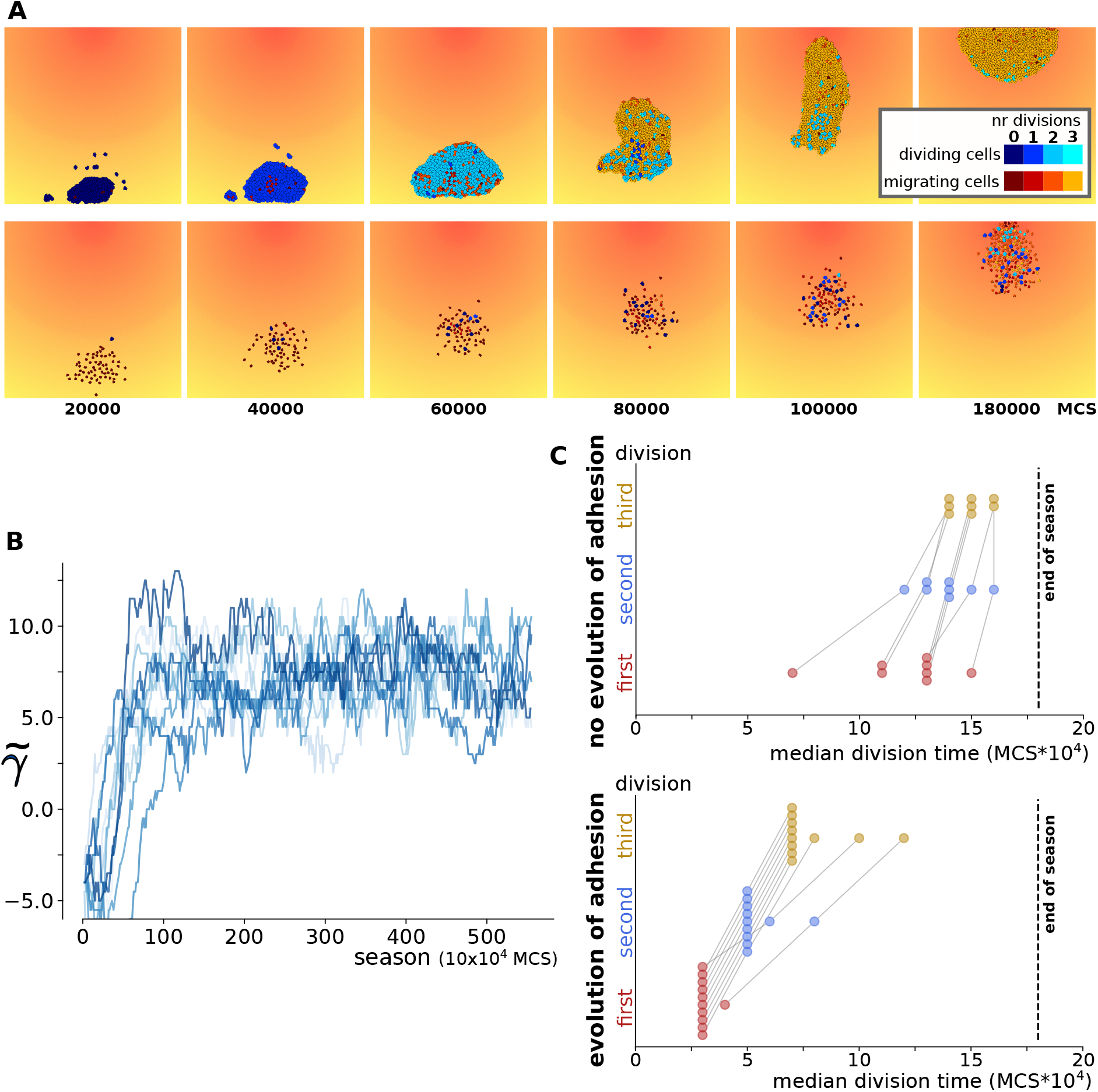
Short season duration and high death rate: either division-late or division-early evolve depending on whether adhesion co-evolves. **a)** Snapshots of the population over the course of one season, in two simulations where different regulatory strategies evolved; one with and one without evolution of adhesion (season length 180000 MCS). Colours denote the state of the cell (dividing=blue, migrating=red/yellow; lighter colours indicate larger number of cell divisions). **b)** Evolution of adhesion in 15 independent simulations with high death rate and short seasons. A greater median *γ* (calculated from the interfacial energy between cells and with the medium, see Methods) indicates stronger adhesion between cells. **c)** The median timing of cell divisions of the last 10 seasons in simulations with short seasons and high death rate, either with or without evolution of adhesion Lines between dots connect values belonging to the same simulations (see also Supp. Fig. S3).

**Table 1:**
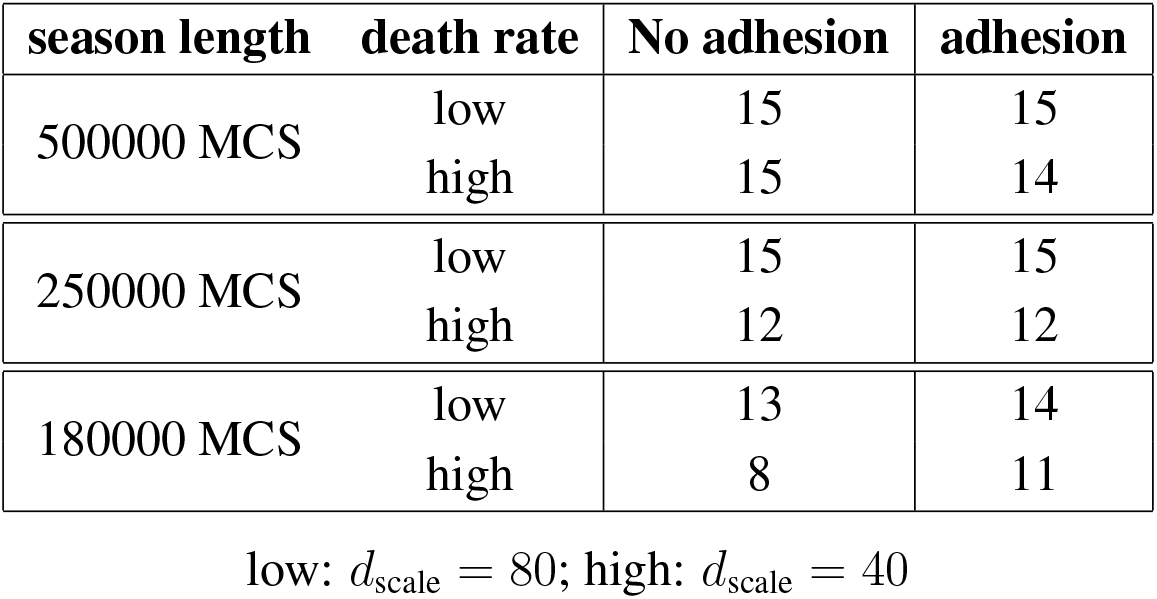
Number of surviving populations (out of 15)

| season length | death rate | No adhesion | adhesion |
| --- | --- | --- | --- |
| 500000 MCS | low | 15 | 15 |
|  | high | 15 | 14 |
| 250000 MCS | low | 15 | 15 |
|  | high | 12 | 12 |
| 180000 MCS | low | 13 | 14 |
|  | high | 8 | 11 |

low: $d_{\text{scale}} = 80$ ; high: $d_{\text{scale}} = 40$

**Table 2:**
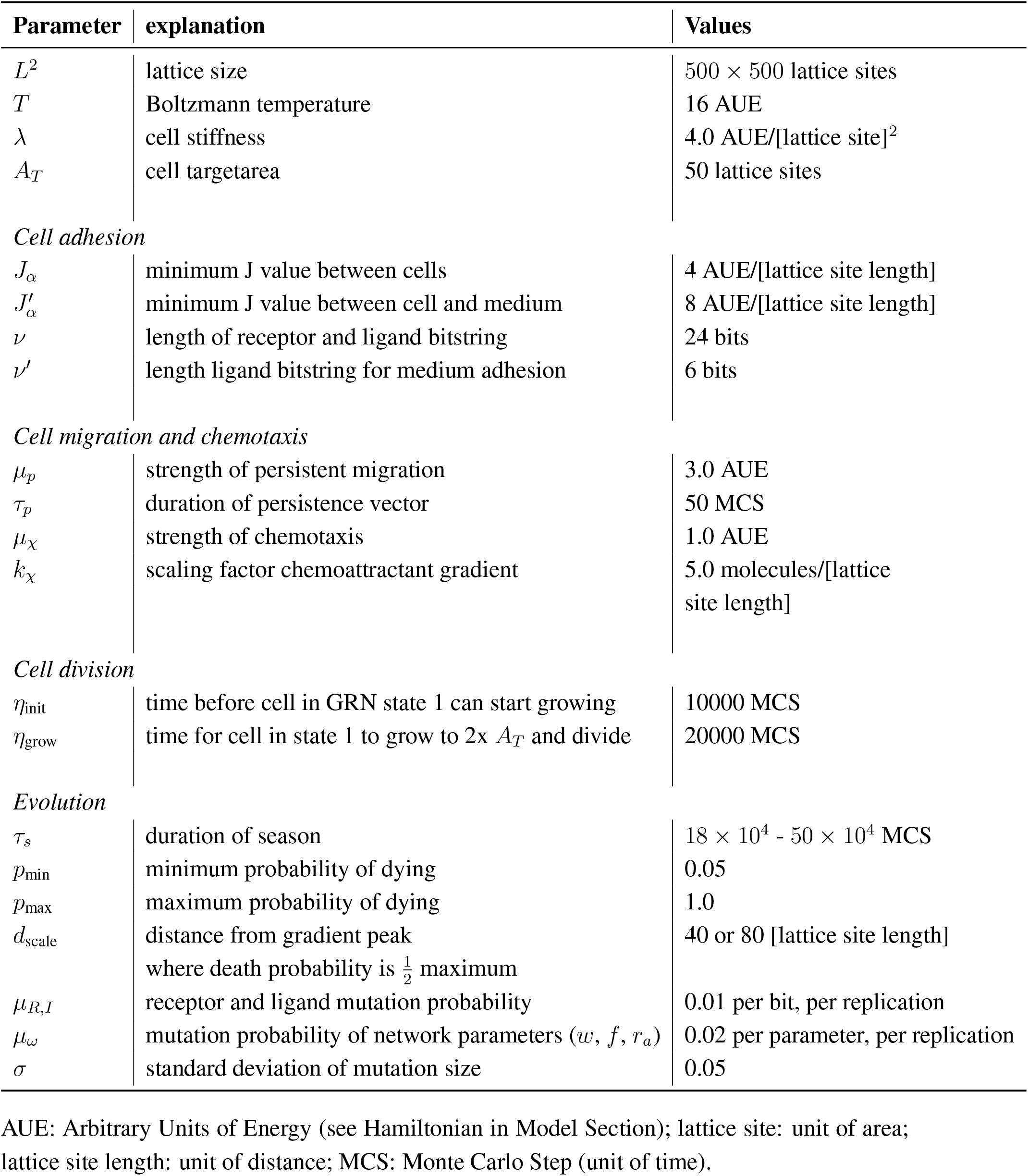
Parameters.

| Parameter | explanation | Values |
| --- | --- | --- |
| $L^2$ | lattice size | $500 \times 500$ lattice sites |
| $T$ | Boltzmann temperature | 16 AUE |
| $\lambda$ | cell stiffness | $4.0 \text{ AUE}/[\text{lattice site}]^2$ |
| $A_T$ | cell target area | 50 lattice sites |
| *Cell adhesion* |  |  |
| $J_\alpha$ | minimum J value between cells | 4 AUE/[lattice site length] |
| $J'_\alpha$ | minimum J value between cell and medium | 8 AUE/[lattice site length] |
| $\nu$ | length of receptor and ligand bitstring | 24 bits |
| $\nu'$ | length ligand bitstring for medium adhesion | 6 bits |
| *Cell migration and chemotaxis* |  |  |
| $\mu_p$ | strength of persistent migration | 3.0 AUE |
| $\tau_p$ | duration of persistence vector | 50 MCS |
| $\mu_\chi$ | strength of chemotaxis | 1.0 AUE |
| $k_\chi$ | scaling factor chemoattractant gradient | $5.0 \text{ molecules}/[\text{lattice site length}]$ |
| *Cell division* |  |  |
| $\eta_{\text{init}}$ | time before cell in GRN state 1 can start growing | 10000 MCS |
| $\eta_{\text{grow}}$ | time for cell in state 1 to grow to $2 \times A_T$ and divide | 20000 MCS |
| *Evolution* |  |  |
| $\tau_s$ | duration of season | $18 \times 10^4 - 50 \times 10^4$ MCS |
| $p_{\text{min}}$ | minimum probability of dying | 0.05 |
| $p_{\text{max}}$ | maximum probability of dying | 1.0 |
| $d_{\text{scale}}$ | distance from gradient peak<br>where death probability is $\frac{1}{2}$ maximum | 40 or 80 [lattice site length] |
| $\mu_{R,I}$ | receptor and ligand mutation probability | 0.01 per bit, per replication |
| $\mu_\omega$ | mutation probability of network parameters ($w, f, r_a$) | 0.02 per parameter, per replication |
| $\sigma$ | standard deviation of mutation size | 0.05 |

AUE: Arbitrary Units of Energy (see Hamiltonian in Model Section); lattice site: unit of area; lattice site length: unit of distance; MCS: Monte Carlo Step (unit of time).

We found that a spectrum of strategies evolved in the different simulations, which could be distinguished by the timing of their divisions: early in the season, late, or somewhere in between. When cells divided very early in the season (a “division-early strategy”), they typically performed all three divisions first and then switched to the migratory state (Fig. 2A, top row; Supp. fig. S1B; Supp. Vid. 1,2). In “division-late strategies”, cell behaviour more often depended on where the cell was located on the lattice. Cells typically migrated until they reached a particular concentration of the chemoattractant, performed a division, and then migrated further before dividing again at a higher chemoattractant concentration (Fig. 2A, bottom row; Supp. Fig. S1C; Supp. Vid. 3,4). Intermediate strategies involved cells migrating a short distance before performing the first division. Populations with a more division-late strategy remained smaller compared to division-early strategies, because fewer cells completed all three divisions during the season (Supp. Fig. S2).

For mild conditions (longer seasons, lower death rates) a broader range of strategies evolved in the different simulations, regardless of the presence or absence of adhesion evolution (Supp. Fig. S3). Simulations with shorter seasons (250000 or 180000 MCS) and higher death rate (40) led to progressively more polarised strategies. In simulations where adhesion could not evolve, we found more “division-late” solutions (Fig. 2C, top; Supp Fig. S3), while in simulations where the receptors and ligands for adhesion were allowed to evolve, we found more “division-early” solutions (Fig. 2C, bottom). In the latter simulations, the population also rapidly evolved adhesion and became multicellular (Fig 2B), because of the increased efficiency of collective migration, and the competitive advantage of adhering cells at the peak of the gradient [23]. This suggests that the evolution of adhesion strongly affects the selection on regulatory strategy, particular under high selection pressure.

### Multicellularity selects for the opposite regulatory strategy from unicellular organisms

Under harsh conditions, cell populations that could evolve adhesion, concomitantly evolved the opposite regulatory strategy from populations that remained unicellular. We hypothesise that, as adhesion evolves, cells in the nascent multicellular group experience a novel selection pressure stemming from the new biotic environment, that results in a division-early strategy. Similarly, a division-late strategy might evolve as a consequence of loss of adhesion, when cells revert to a unicellular state. To test this hypothesis, we assess the evolutionary stability of the two regulatory strategies - division-early and division-late - under the opposite adhesion regime.

We selected 4 individuals from different high-death rate, short-season simulations that were evolved without adhesion and let them evolve their adhesion strength. 4 individuals evolved with adhesion were used for simulations without adhesion (*γ* = —4). Each of these individuals was used to start a new population in 5 independent simulations.

All 20 populations that could evolve adhesion, rapidly did so (Supp. Fig. S4). Concurrently, they also evolved a division-early strategy, especially for their first division (Fig. 3A,B). Two factors contributed to earlier divisions: collective migration and the evolved regulation. Collective cell migration speeds up the group’s chemotaxis compared to a single cell [23], allowing cells to reach the chemoattractant concentration at which they divide at an earlier time – particularly when cells have a division-late strategy (Supp. Fig. S6). Their gene regulation also evolved so that divisions occurred at a lower concentration of the chemoattractant gradient (Fig. 3C, Supp. Fig. S5A).

**Figure 3:**
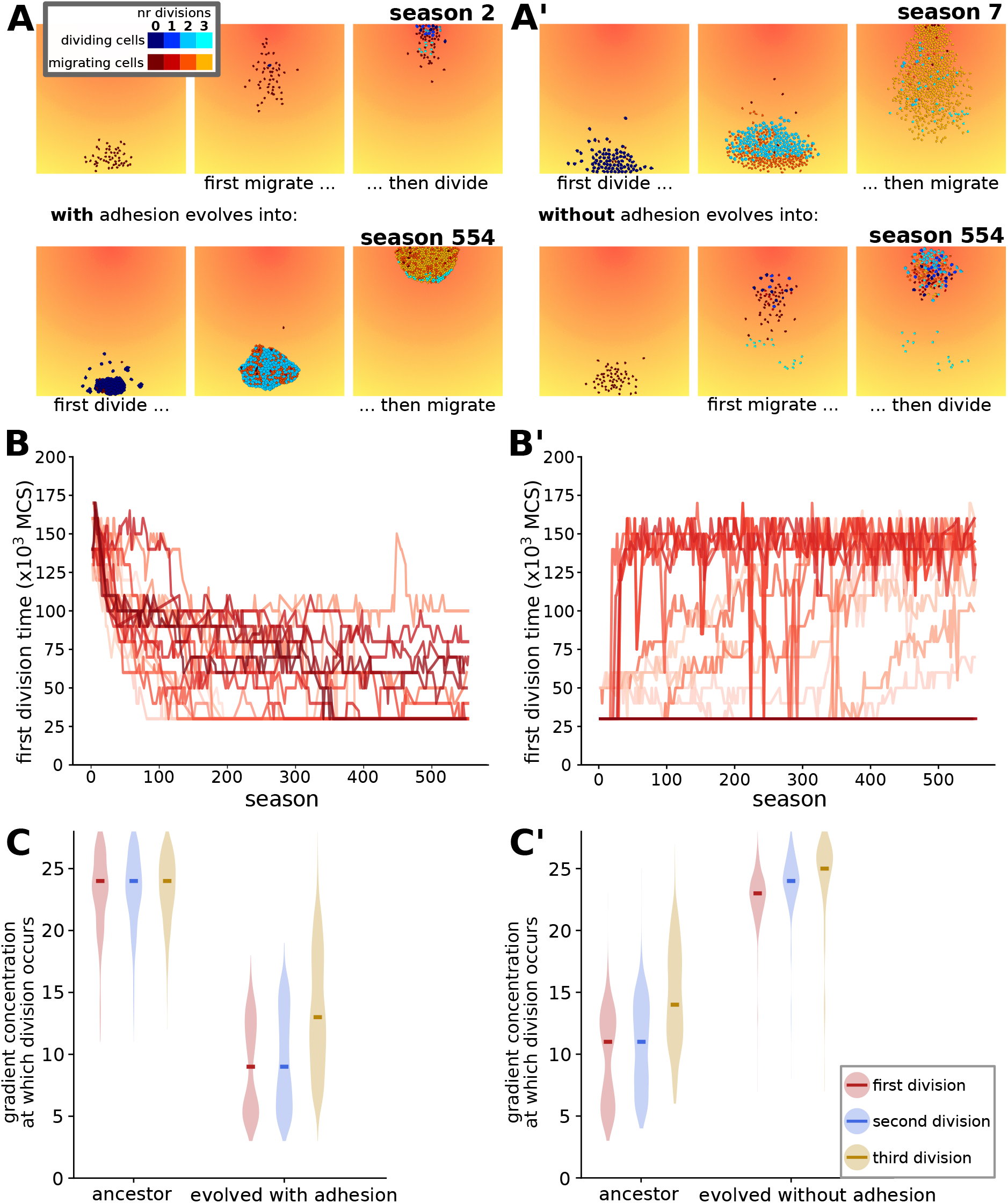
When the adhesion capacity of a population is switched, regulation evolved to the opposite strategy. **a-a’)** Snapshots of a simulation from a simulation with a division-late (a) or a division-early (a’) ancestor that evolved towards the opposite strategy due to a switch in adhesion possibility (a: switch from simulation without adhesion to simulation with adhesion; a’ vice versa.) (seasons chosen where the peak is on the opposite side of the lattice). **b-b’)** The evolution of the timing of the first division, for all simulations with switched adhesion. Median value of the population at any time point. Colour distinguishes the different simulations in one plot. **c-c’)** The concentration at which the three divisions occur, for an ancestor and descendant populations in 1 of the 5 replicate simulations. The thick lines indicate the median of the population. All other replicates can be found in Supp. Fig. S5.

Instead, the individuals that were switched to simulations without adhesion gave rise to populations which evolved towards a division-late strategy Fig. 3A’, B’). The main cause for this was a change in regulation: cells evolved to divide at a higher chemoattractant concentration (Fig. 3C’; Supp. Fig. S5B). We also observed that non adhering cells with a division-late strategy physically hindered each other while moving towards the gradient, leading to a slight delay in reaching the signal to divide compared to single cells (Supp. Fig. S6). Although this effect was small, it contributed to the overall competition between cells in a division-late strategy.

In summary, the evolutionary simulations show a clear change in regulatory strategy in reaction to a change in adhesion (Fig. 4A-B). The speed of adaptation however depended on the ancestral strategy, with the division-late strategies typically being faster to change to a division-early strategy than vice versa (compare Fig. 3B with B’). These experiments recapitulate the evolutionary transition from a unicellular ancestor to a multicellular group, and the evolutionary reversal from a multicellular to a unicellular state. Taken together, they suggest that transitioning towards multicellularity is evolutionarily easier than its reversal (Fig. 4C).

**Figure 4:**
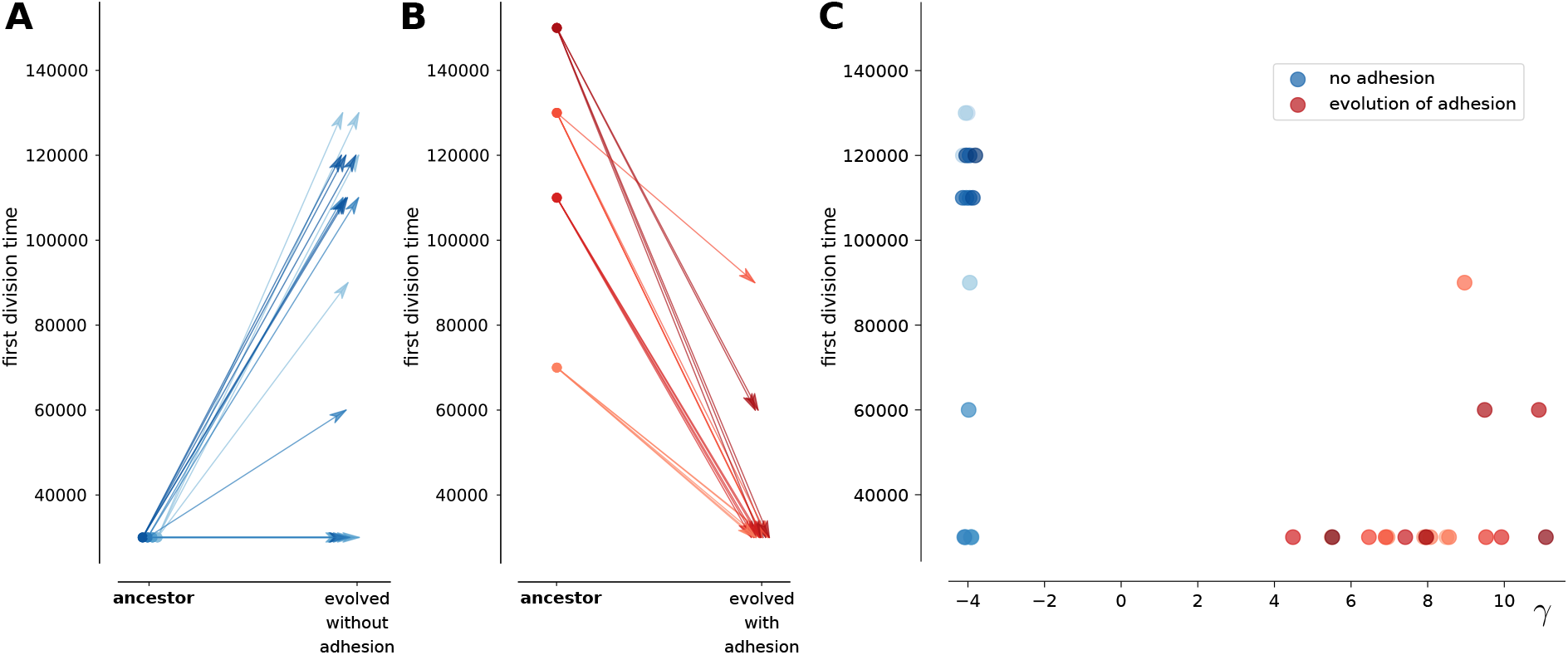
Summary of changes in division time in all simulations. **a-b)** Average timing of the first cell division in the last ten seasons of the simulations **a)** evolved without cell adhesion **b)** the possibility of evolving adhesion. The left points in the graph indicate the division timing the ancestral strategy with which continued simulations were started. Four different ancestral strategies from simulations without adhesion were allowed to continue evolution with adhesion, and vice versa (5 independent simulations per ancestral strategy). **c)** Comparison of evolved adhesion and division timing after the switch. For all simulations evolved without adhesion, *γ* is fixed to −4 (jitter added in plot for clarity).

### Adhesion drives selection for selfish early divisions by lifting pressure to reach the peak

We next analyse the competition and cooperation dynamics during the transition to multicellularity in our model. We start from a division-early ancestor that is evolving cell adhesion (as in Fig. 3A), and consider two regulatory mutants: one that migrates for longer before dividing, and one that divides earlier. The first mutant could be considered more cooperative, because it foregoes replication to carry the group. The second then could be considered more selfish, because it gets carried by the group while prioritising its own replication. We consistently observe that the second strategy takes over, suggesting that it evolves due to competition within the cluster.

We set up competition experiments to investigate how population dynamics cause a selective advantage for division-late strategies in unicellular populations, and for division-early strategies in multicellular groups. We placed two groups of cells with different strategies and/or different adhesion strengths next to each other at the end of the lattice opposite the peak of the gradient (Fig. 5A). Cells were allowed to migrate and divide, but no mutations could happen upon division, so that all cells in a group kept the same strategy. Cell death occurred at the end of each season as in the evolutionary simulations, and simulations were run until one of the groups went extinct.

**Figure 5:**
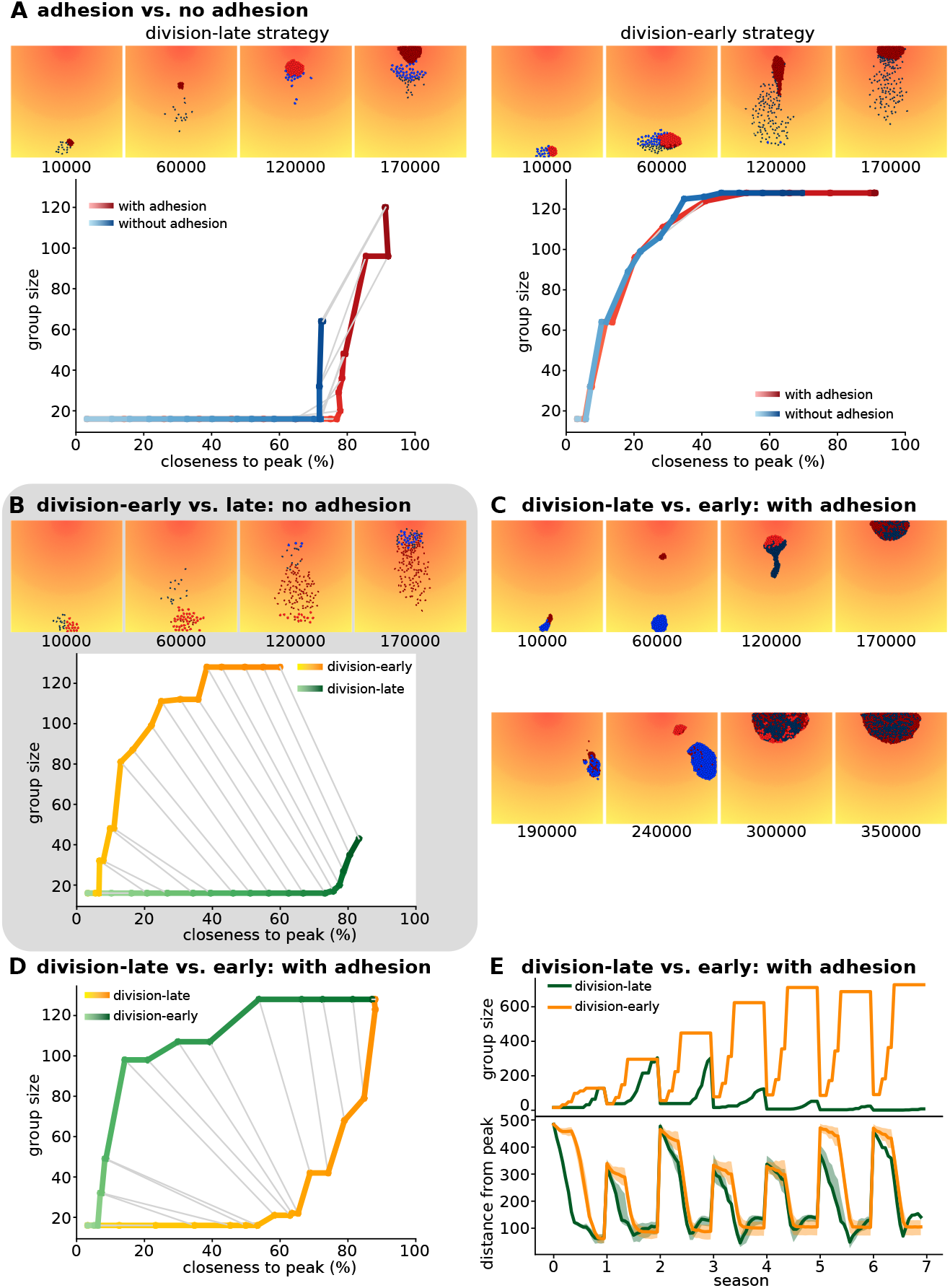
Competition experiments show that selective advantage of a regulatory strategy depends on presence of adhesion. Setup of competition experiment: two groups of each 16 cells are placed adjacent to each other, with differing regulatory networks or differing adhesion values. The simulation is then run for multiple season (without mutations), until one of the groups has gone extinct. The cells of one group are coloured red (light red when dividing) and the others are blue (light blue when dividing) **a-d)** Snapshots show the first season of the competition simulation. Graph plots group size against median distance of cells to the peak for the first season (0%=maximum distance, 100%=at the peak). Light-to-dark colour gradient of the line indicates time in the season, grey lines connect equal time points between the two groups. Repli-cates can be found in Supp. Fig. S7. **a)** Competition between adhering (*γ* = 6, red) and non-adhering (γ = —4, blue) cells, either both with a division-late strategy (left) or division-early strategy (right). **b)** Competition between ancestral, division-early strategy (evolved with adhesion before the switch to non-adhering conditions; red in snapshots) and evolved, division-late strategy (blue in snapshots); both non-adhering. **c)** Two seasons of the competition experiment between two adhering groups, one with an ancestral, division-late strategy (evolved without adhesion before the switch to a simulation in which adhesion could evolve; red in snapshots) and one with a strategy evolved after 500 seasons with possibility of adhesion (having become more division-early; blue in snapshots). **d)** Graph of the first season of the competition experiment in c **e)** Time dynamics of the competition in c. Top plot shows group sizes, bottom plot shows distance to the peak. Shading indicates 25^th^ and 75^th^ percentile of population.

Adhering groups always won against non-adhering groups with the same regulatory strategy (Fig. 5B). We observed that the main advantage to evolving adhesion came from adhering groups displacing non-adhering cells at the peak of the gradient. Collective chemotaxis was beneficial but had a smaller effect (Fig. 5B and Supp. Vid. 5,6). Therefore, adhesion evolves because it increases interference competition.

Then we investigated why the opposite regulatory strategy evolves when the adhesion regime is inverted (from division-early towards division-late when switched from adhesion to no adhesion, and vice versa). We let a division-early and a division-late population compete without adhesion (we set *γ* = −4). In this case, the division-late cells were closer to the peak throughout each season (Fig. 5C). The division-late strategy won because non-adhering cells exclude each other from the peak. The cells that reached the peak sooner were locked in place by the pressure exercised by the later-arriving migratory cells. (Supp. Vid. 7). However, in the first season there were significantly fewer division-late cells, indicating that this strategy sacrificed cell divisions to get to the peak sooner.

We then let a division-late group compete with division-late group, assigning both groups a high adhesion value in the competition experiment (*γ_c,m_* = 6). The competition dynamics of these two groups were more complex. The division-late group reached the peak of the gradient earlier than the division-early group. While the latter reached the peak with a larger population size, both groups had the same population size at the end of the season. Because adhesion was mediated by identical ligands and receptors, the two groups mixed freely. However, when we plotted the distance of both groups over multiple seasons, we found that the division-late group did not remain close to the peak until the end of the season, despite arriving there earlier (Fig. 5E). The videos show that the division-late cells were displaced because many still performed divisions close to the peak, and were therefore in a non-migratory state (Supp. Vid. 8). This also resulted in smaller population sizes in later seasons, as the larger mass of the division-early cells kept the division-late cells from reaching high enough concentrations. Thus, as adhesion increases mixing between sub-populations at the peak, a division-early strategy is more competitive because it results in larger numbers of actively motile cells.

In summary, when cells cannot evolve adhesion, their evolution is driven by a competition for reaching and occupying the peak first. This leads to a division-late strategy, where cells combine the use of the abiotic environmental cues (the gradient) with the information on their internal state. This strategy yields a survival benefit due to lower death rates, but at the cost of fewer divisions. When cells can (and do) evolve adhesion, they optimise the use of their biotic environment, i.e., the other cells. Adhesion allows cells to reduce interference competition at the peak of the gradient by increasing cell mixing, and exploit each other for migration while maximising their divisions. Cells therefore become more selfish as adhesion – and thereby collective migration – evolves. The change in selection pressure as the cellular context changes is the hallmark of differentiated multicellular organisation. Finally, the multicellular clusters often displayed spatial patterns of cell differentiation that resembled development, with morphogenetic events such as stretching and compression of the multicellular cluster (reminiscent of convergent extension [29, 31]). This is especially clear for the dynamics of a genetically homogeneous cluster (Fig. 6, Supp. Vid. 9). Unlike in complex embryonic development however, the differentiation in our simulations remains temporary in nature, with cells continually switching between a migratory and dividing state.

**Figure 6:**
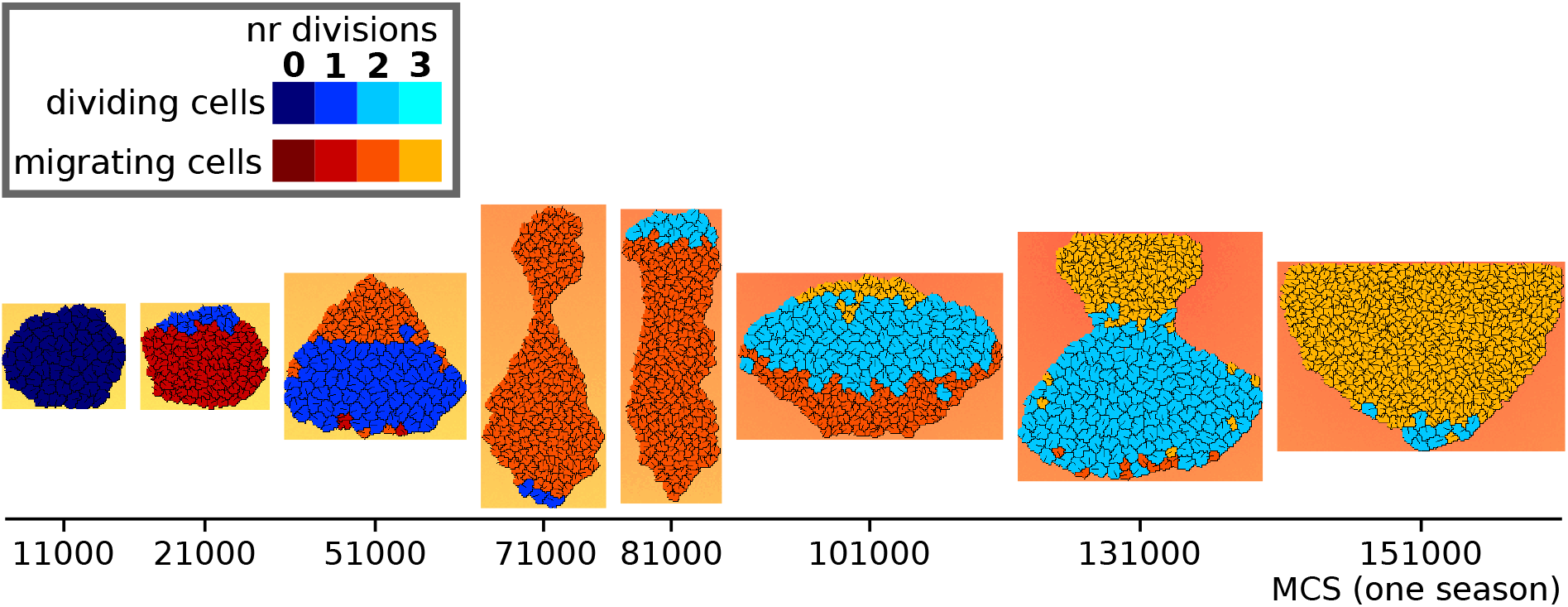
Development of genetically homogeneous cluster. Images of a developing cluster over one season. The season was started with 50 genetically identical cells with a regulatory strategy taken from from a simulation evolved with adhesion (death rate = 80, season duration = 180000). Images are centered on the cluster as it moves through the grid.

## Discussion

In this study, we show how the evolution of adhesion changes the competition dynamics between cells that are able to sense and react to their environment. In both the unicellular and the multicellular case, cells selfishly maximise their own replication or survival, but the evolved strategy to do so depends on the abiotic and cellular context. In the model, cells evolve to regulate when to migrate towards resources and when to divide. Evolution prioritises migration in the unicellular case (when adhesion cannot evolve), because of competition to reach the resources necessary for survival. Adhesion facilitates collective migration and cell mixing, thereby lifting the selection pressure to reach the resources first. In this case, cells compete to perform all divisions while remaining in the migrating cluster, and to remain as close to the peak as possible by maintaining a migratory state once there. Thus, the abiotic environment determines competition in absence of adhesion, while the biotic environment – the other cells – determines competition in the presence of adhesion. We therefore expect that at the transition to multicellularity, the regulatory program of unicellular organisms drastically changed its dynamics to suit the new (cellular) environment – yielding proto-developmental patterning as a side effect of the process.

Current bioinformatic evidence from the holozoan relatives of animals points to a unicellular ancestor that had a complex life cycle, possibly with a multicellular stage, and a significant regulatory toolkit [10]. In fact, many of the genes necessary for coordinating multicellular development, including adhesion proteins, transcription factors and signalling genes, were already partially present in the unicellular ancestor and extant closely related unicellular species [8, 9, 32–35]. Our results show that the ancestral toolkit could have undergone rapid evolution to yield new strategies for competition within a newly multicellular context. We find that the temporal behaviour of cells, which results from their individual decoding of the environment, can already yield transient spatial patterning in the multicellular cluster. The pattern is further stabilized by the differential sorting between migrating and dividing cells. No large changes to the available genetic toolkit are required for such transient patterning, only refinements of the existing regulation.

Cell states are organised by a gradient produced by the embryo in complex multicellular organisms. In contrast, external and internal cues combine to generate pattern formation in simpler multicellular organisms. For instance, in the multicellular alga *Volvox carteri,* dark-light transitions govern the differentiation of cells into somatic and germline cells, depending on their size after embryogenesis [36]. Similarly, in the multicellular (slug) phase of the slime mould *Dictyostelium discoideum’s* life cycle, cells migrate up external gradients of heat and light, causing different cell types to sort into specific regions along the body of the slug [37, 38]. Environmental cues triggering multicellularity can also be biotic: bacterial molecules induce rosette formation or swarming in the choanoflagellate *Salpingoeca rosetta* [39, 40], while some myxobacteria [41] and slime moulds migrate collectively to feed on bacteria [42]. Previous models have highlighted the effect of motility on the evolution of adhesion in social [43, 44] and non-social groups [23]. Our current model suggests that the transition to multicellularity may be driven by collective migration and increased competition within the cell cluster, and that external gradients may have provided the cues for spatial pattern formation and cell differentiation. Experimental evolution with unicellular or transiently multicellular phototactic or chemotactic species may provide a test of these predictions.

We do not observe a complete temporal-to-spatial shift in these simulations, as has been hypothesized for the transition to multicellularity [45]. Cells keep switching between states to maximise their own growth and survival. We expect that multilevel competition between multiple distinct multicellular clusters may be required to select for stable differentiation, as division of labour between cells may result in more successful groups. The ensuing group-level competition may also allow cooperative strategies to arise. In this respect, our work shows that the transition to multicellularity does not have to be accompanied by cooperation between cells. Cooperation in the multicellular group can be a later innovation, perhaps under the control of a germ-line. Additionally, the evolution of stable patterning may require a more evolvable genetic toolkit, which can expand to facilitate more complex regulation of cell-state, adhesion and cell-cell signalling. Future work may then be able to assess how the genetic toolkit used by the unicellular ancestor to regulate cell behaviour, formed the basis for stable cell differentiation and division of labour in multicellular clusters – as has been suggested before [3, 11, 12].

## Conclusions

In our previous study [23], we found that adhesion can evolve in response to an emergent selection pressure for collective migration in a noisy environment. Here, we find that such evolution of adhesion strongly determines the competition dynamics between cells, pushing it from a competition for survival to a competition for reproductive success. Adhesion therefore changes the evolution of behaviour regulation. In combination with an external chemoattractant gradient, this leads to the emergence of differentiation patterns in newly multicellular organisms: selfish multicellularity.

## Declarations

### Ethics approval and consent to participate

Not applicable.

### Consent for publication

Not applicable.

### Availability of data and materials

The source code used for the simulations in this study is freely available from Github: [46]. The simulation data that were generated for the current study are available from the corresponding author on reasonable request.

### Authors’ contributions

RV and ESC jointly conceived and designed the study, contributed to code development, analysed the simulation outcomes and contributed to the writing of the manuscript. RV wrote the scripts for data analysis and wrote the first draft. Both authors read and approved the submitted version.

## Acknowledgements

We thank Bram Hoogland and Pjotr van der Jagt for their thorough reading of the draft and helpful suggestions.

## List of abbreviations

CPM: Cellular Potts Model
MCS: Monte Carlo Step
GRN: Gene regulatory network

## Methods

We model an evolving population of cells that can adhere to each other, divide, migrate and perform chemotaxis on a two-dimensional lattice containing a gradient of chemotactic signal. Cells consist of multiple lattice sites, giving them an explicit shape and volume. They also contain an evolvable gene regulatory network that senses the environment and regulates the decision to either divide or migrate (explained below). Cell dynamics on the lattice (movement, adhesion) are governed by the Cellular Potts Model (CPM) formalism [24, 25] and simulated with a Monte Carlo method. The population undergoes a seasonal culling, in which cells further from the gradient source have a higher chance of dying. Then, the gradient source is placed at a different position, after which the new season starts. The custom software, written in C++, can be found at [46].

### Regulation of cell behaviour

Cells have a simple evolvable regulatory network that determines their behaviour: either migrating and following chemotactic signals, or growing and dividing. The overall network architecture has been previously used to model gene regulation in micro-organisms [30], and consists of 2 sensory nodes, 3 regulatory nodes and an output node, totalling *N* = 6 nodes (Fig. 1B). The sensory nodes receive the local concentration of chemoattractant (a real number between 0 and 28) and the number of times the cell has already divided. The regulatory nodes receive input from the sensory nodes and the other regulatory nodes. The output node takes as input the state of the regulatory nodes, to determine whether the cell is in a migratory or dividing state. The regulatory nodes and output node are Boolean. Their new states are calculated synchronously from the previous state, as follows:

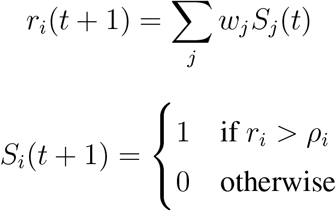

where *r_i_*(*t* + 1) is the regulatory input into node *i* from nodes *j* in state *S_j_* at time *t*, which regulate *i* with weight *W_j_*. *S_i_*(*t* + 1) is then the new state of node *i*, at time *t* +1. The new state is 1 if the value of the regulatory input *r_i_* is greater than its activation threshold *ρ_i_*. When the node *j* is an input node, *S_j_* is the value of the input node times an evolvable scaling factor *φ_j_*; otherwise it is the Boolean state of a regulatory node (including *i* itself). Since the update is synchronous, the state of all nodes is only updated after all new node states are calculated from the old node states. For each cell, the network state is updated every 20 MCS, to even out microscopic cell fluctuations. The networks of different cells are updated at different MCS to prevent artificial synchrony between cells.

When the state of the output node is 0, the cell is in migratory mode. When the state is 1 for *η*_init_ consecutive timesteps, the cell enters the dividing cell state, during which it does not migrate or perform chemotaxis. It will need *η*_grow_ timesteps to grow to twice the size (by increasing target size, *A_T_*, every *η*_grow_/*A_T_* MCS) and prepare for division before it can actually divide. If, in that time, the output node becomes 0, the cell will shrink again and revert back to the migratory state, and will start the division process anew if the output node becomes 1 again. Once the cell divides, the state of all nodes is reset to 0. A cell can divide a maximum of three times per season; when the output node remains one after that, the cell is non-migratory, but won’t divide again.

Upon cell division, the regulatory network is passed to the daughter cell with the possibility of mutations. Each parameter of the network (*w_i_, φ_i_, ρ_i_*) mutates independently during cell division. Mutations occur with probability *μ_ω_* and change the values of these parameters by a small random number sampled from a normal distribution with mean 0 and *σ* = 0.05.

### Cellular Potts Model

The Cellular Potts Model dynamics are implemented with the Tissue Simulation Toolkit [26], on a regular square lattice 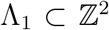 of size *L* × *L*. A cell *c* consists of the set of lattice sites 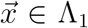 with the same spin *s*, i.e. 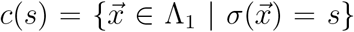. The chemotactic signal is located on a second plane Λ_2_, of the same size and spacing as Λ_1_.

The cells move in the Λ**1** lattice due to the displacement of their boundary, arising from stochastic fluctuations. These fluctuations tend to minimise a cell’s energy, whose terms correspond to biophysically motivated cell properties ([47]). The Λ**1** lattice is updated through the Metropolis algorithm. Each Monte Carlo step (MCS), *L* × *L* lattice sites are drawn randomly. For each site belonging to the boundary of a cell, a random site from its Moore neighbourhood, 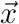, is selected which may copy its spin value 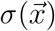 into this lattice site. To speed up simulations, we used an algorithm that only considers updates for neighbouring lattice sites that have a different s, as implemented in [48]. Whether an attempted spin copy is accepted depends on the contribution of several terms to the energy *H* of the system, described by the Hamiltonian, as well as other biases *Y*. A copy is always accepted if energy is dissipated, i.e. if Δ*H* + *Y* < 0 (with Δ*H* = *H*_after copy_ — *H*_before copy_), and may be accepted if Δ*H* + *Y* ≥ 0 because of “thermal” fluctuations that follow a Boltzmann distribution:

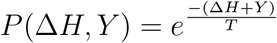

with *T* =16 the Boltzmann temperature (in Arbitrary Units of Energy AUE), which controls the probability of energetically unfavourable copy events. The Hamiltonian consists of two terms, corresponding to cell size maintenance and adhesion:

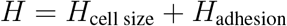

The copy biases, or “work terms”, *Y* consist of terms corresponding to cell migration and chemotaxis:

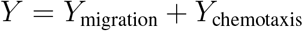

### Cell size maintenance

Cell size *A*(*c*) = |*c*(*s*)|, the number of lattice sites that compose a cell, is assumed to remain close to a target size *A_T_*. Deviations from the target size are inhibited by adding the following term to the Hamiltonian:

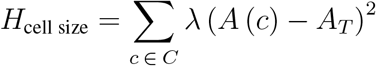

with *C* the set of cells and λ representing the resistance of cells against volume changes. A cell’s target volume may grow when it prepares to divide, and is halved again upon division (see also below).

### Cell adhesion

The total adhesion energy resulting from interfaces between cells and with the medium is implemented as:

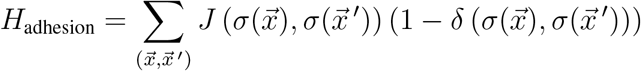

summing over all pairs of neighbouring pixels with different as: 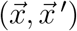. 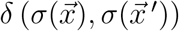 is the Kronecker delta which restricts the energy calculations to the interfaces.

As previously described [23], cells express ligand and receptor proteins on their surface that determine the surface energy of cell interfaces, 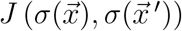. Ligands and receptors are modelled as binary strings of fixed length *ν* (Fig. 1). Cell adhesion increases (i.e. lower *J* values) with greater complementarity between their receptors *R* and ligands *I* (i.e. larger Hamming distance 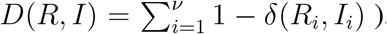, where *R_i_* and *I_i_* are corresponding bits in the receptor of one cell and the ligand of the other cell, and *δ* is the Kronecker delta function which is 1 when the two bits match. Thus, given two cells with spin values *σ*_1_ and *σ*_2_ and their corresponding pairs of receptors and ligands (*R*(*σ*_1_), *I*(*σ*_1_)) and (*R*(*σ*_2_), *I*(*σ*_2_)):

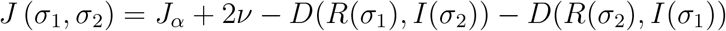

with *J_α_* = 4 so that *J* (*σ*_1_, *σ*_2_) ranges between [4, 52].

Adhesion of a cell with medium is assumed to depend only on the cell (the medium is inert, i.e. *J* (*σ*_medium_, *σ*_medium_) = 0), and in particular it depends only on a subset of the ligand proteins of a cell. This subset consists of the substring of *I, I*_[0;*<*]_, which begins at the initial position of *I* and has length *ν*’ The value of *J* (*σ*_1_, *σ*_medium_) is calculated as:

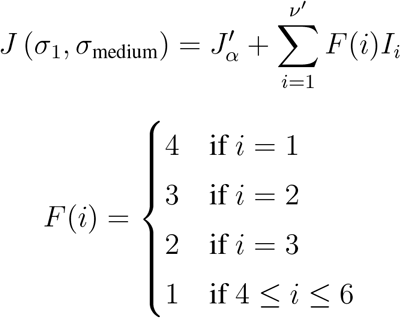

with 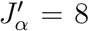 and *F*(*i*) a piece-wise defined function (a lookup table). Thus, each bit *I_i_* in *I*_[0,*ν*’]_ contributes to *J* (*σ*_1_,*σ*_medium_) with a different weight, resulting in *J* values falling in the interval [8, 20].

The strength with which cells adhere to each other depends on both the J value between the cell and the medium, and the J value of the interaction between cells. This can be represented by the *γ* value, the surface tension between cells and medium:

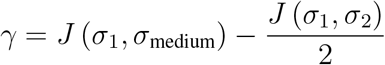

When *γ* is negative, cells preferentially interact with the medium and do not adhere. In simulations where adhesion cannot evolve, we set *J* (*σ*_1_, *σ*_2_) = 36 and *J* (*σ*_1_, *σ*_medium_) = 14 → *γ* = −4, yielding nonadhering cells. To simplify the notation in the main text, we use the subscripts *c* and *m* to refer to lattice sites belonging to cells and medium, so that *J_c,m_* = *J*(*σ*_1_, *σ*_medium_) and *J*_c,c_ = *J*(*σ*_1_, *σ*_2_), where the subscript c,c always assumes two different cells.

In a subset of simulations, the bitstrings of the receptor and ligand responsible for adhesion are mutated upon cell division. These mutations occur in only one of the two daughter cells (chosen at random) with a per-position probability *μ*_R,I_. Mutations flip individual bits (from 0 to 1, and vice versa).

### Cell migration

We model migration (following [49]) by biasing cell movement to their previous direction of motion 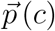: extensions of a cell are energetically more favourable when they are closer to the direction of that cell’s 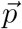:

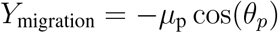

Where *μ*_p_ is the maximum energy contribution due to migration, and *θ_p_* is the angle between 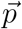 and the vector that extends from the center of mass of the cell to the lattice site into which copying is attempted. Every *τ_p_* MCS the vector 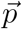 is updated to reflect the actual direction of displacement of the cell over the past *τ_p_* MCS (scaled to unit) (Fig. 1). We set *τ_p_* = 50 MCS. Whether a cell migrates depends on its internal state. Dividing cells do not migrate, so their *μ_p_* = 0. Non-dividing cells have *μ*_p_ = 3. Note that all cells have the same *τ_p_,* but they update their vectors at different MCS to prevent them from moving synchronously.

### Chemotaxis

Individual cells are able to migrate towards the perceived direction of a chemoattractant gradient. In contrast to [23], the slope of the gradient is steeper and we removed a source of noise in the gradient signal, allowing individual cells to identify the location of the peak with ease.

The chemotactic signal is represented by a collection of integer values on a second two dimensional lattice (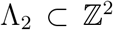, with the same dimensions as the CPM lattice), which remain constant for the duration of one season (*τ_s_* MCS). The amount of chemotactic signal *χ* is largest at the peak, which is located at the center of one of the lattice boundaries, and from there decays linearly in all directions, forming a gradient: 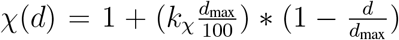, where *k_χ_* is a scaling constant, *d* is the Euclidean distance of a lattice site from the peak of the gradient, and *d*_max_ is the maximum distance between the source of the gradient and any lattice site in Λ_2_. Non integer values of *χ* are changed to ⌈*χ*⌉ (the smallest integer larger than *χ*) with probability equal to ⌈*χ*⌉ – *χ*, otherwise they are truncated to ⌊*χ*⌋ (the largest integer smaller than χ).

A cell s only perceives the chemotactic signal on the portion of Λ_2_ corresponding to sites on Λ_1_ with that spin. We define the vector 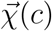 as the vector that spans from the cell’s center of mass to the center of mass of the perceived gradient. Copies of lattice sites are favoured when they align with the direction of the vector 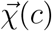, i.e. when there is a small angle *θ_c_* between 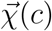 and the vector that spans from the center of mass of the cell to the lattice site into which copying is attempted (Fig. 1):

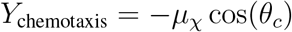

where *μ_χ_* is the maximal propensity to move along the perceived gradient, and is set to *μ_χ_* =1 except for dividing cells, whose *μ_χ_* = 0. A uniform random *θ_c_* ∈ [0, 2*π*[is chosen whenever 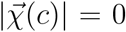, i.e. when, locally, there is no gradient (due to the chemoattractant amount at each lattice site being rounded to integer values).

### Seasonal dynamics

A population of *N* cells undergoes the cell dynamics described above for the duration of a season, i.e. *τ_s_* MCS. Cells can divide throughout the season, and upon division will pass on their regulatory network to the daughter cell with the possibility of mutations (described above). At the end of the season, we assess the distance of each cell from the peak of the chemokine gradient. The further the cell, the higher its probability of being killed before the new season starts. This is implemented as:

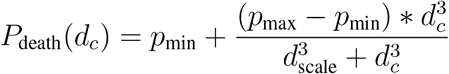

where *p*_min_ is the minimum probability of dying, *p*_max_ the maximum probability, *d_c_* is the distance of cell *c* to the peak of the gradient, and *d*_scale_ is the distance at which the death probability is at half of the maximum. The smaller *d*_scale_, the more cells will die.

## Supplementary material

**Figure S1:**
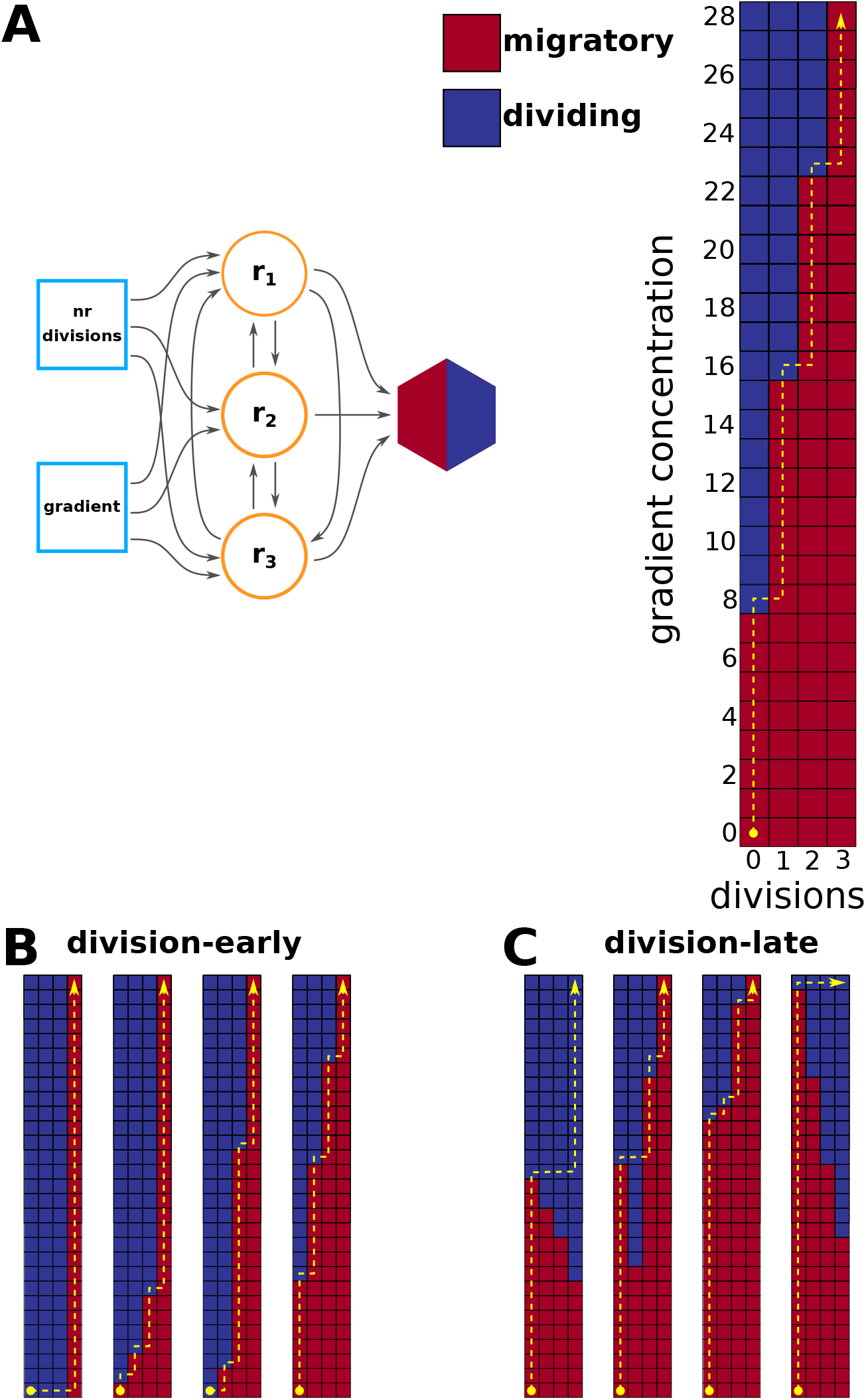
Assessment of evolved GRN responses in unicellular simulations. **A)** We assess the steady state of the output node for all combinations of values for the input nodes (gradient concentration and number of divisions) that are possible in the simulation (with the other nodes starting at 0), and then assign that combination a color in the 2D profile, with red=migratory (output node 0), blue=dividing (output node 1). The profile can be read by starting on the bottom-left, moving up when the pixel is red (migrating to higher concentration), and right when it is blue (dividing and increasing the number of divisions the cell has done). **B)** Examples of GRN responses of division-early, mixed, and division-late strategies. These have then been used for the simulations with harsher conditions.

**Figure S2:**
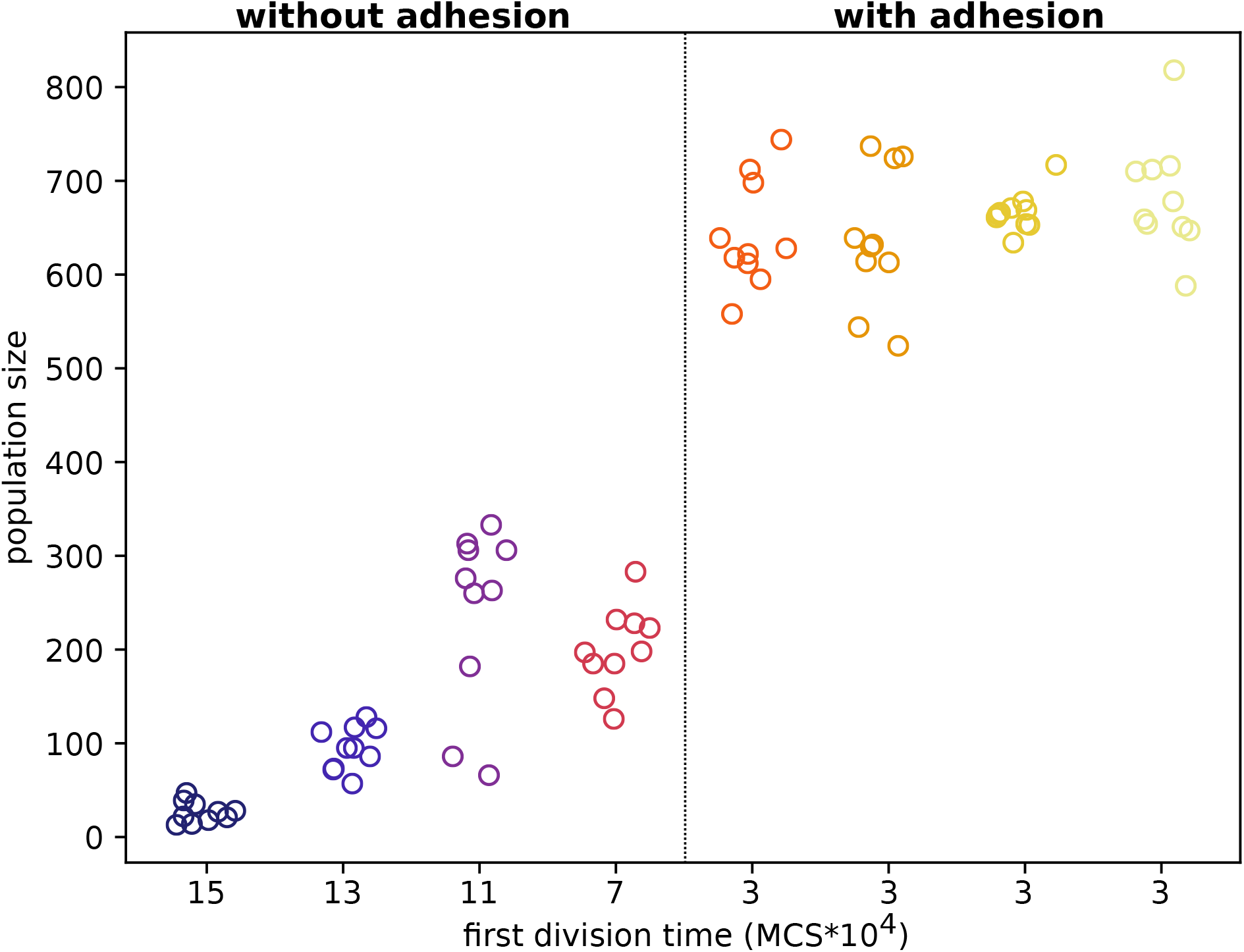
Evolved population size. The population sizes at the end of the last ten seasons in simulations with high death rate and very short seasons. Simulations are sorted by their median first division times (indicated on the x axis), from the simulations that were then used to pick ancestors for the switched simulations. Colours group data from the same simulation. Resolution on division timing was limited to 10000 MCS.

**Figure S3:**
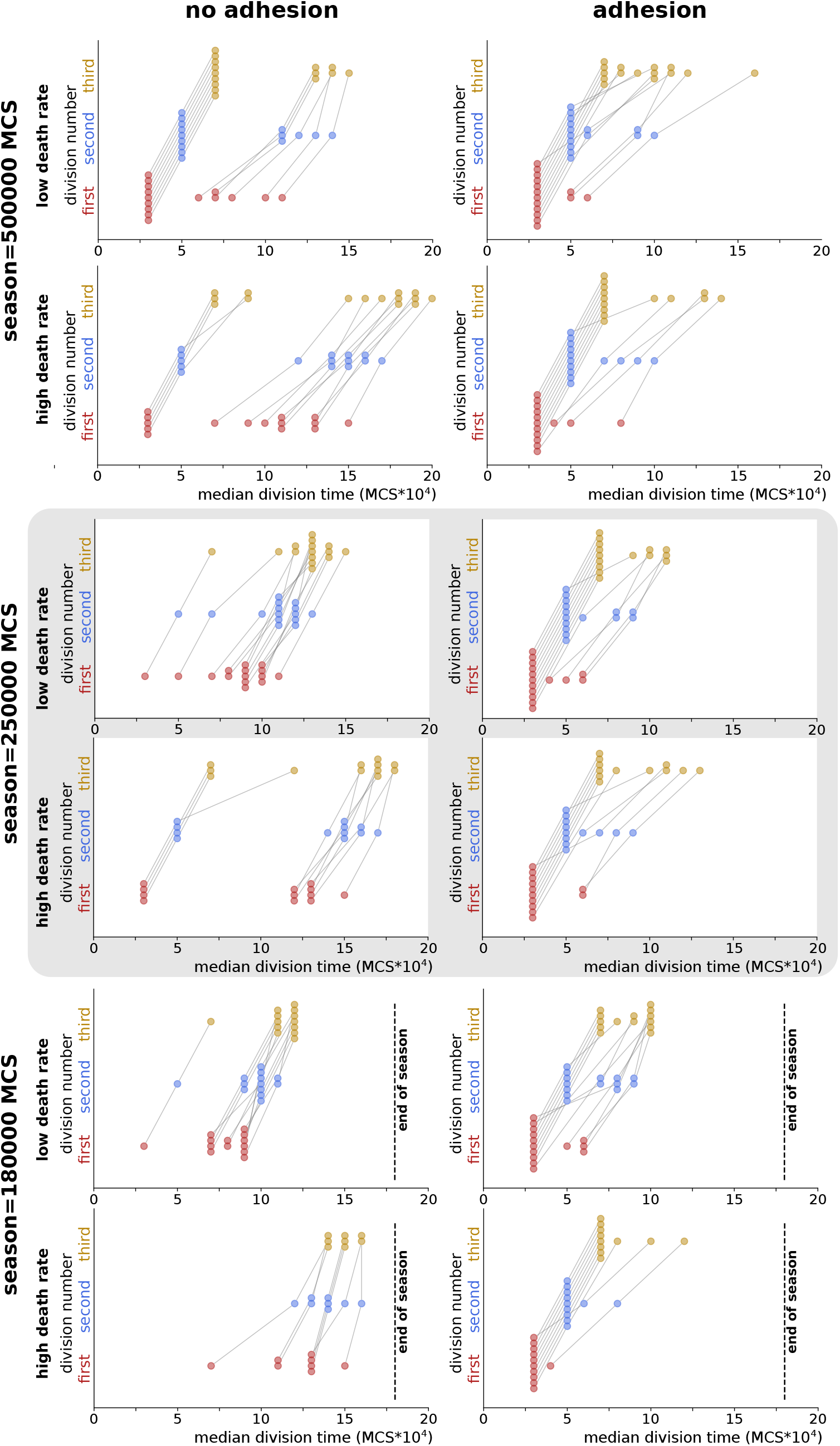
Evolved division timing strategies in all simulation sets. The median timing of cell divisions of the last 10 seasons in all sets of simulations, with varying season duration, death rates and with or without evolution of adhesion. Lines between dots connect values belonging to the same simulations.

**Figure S4:**
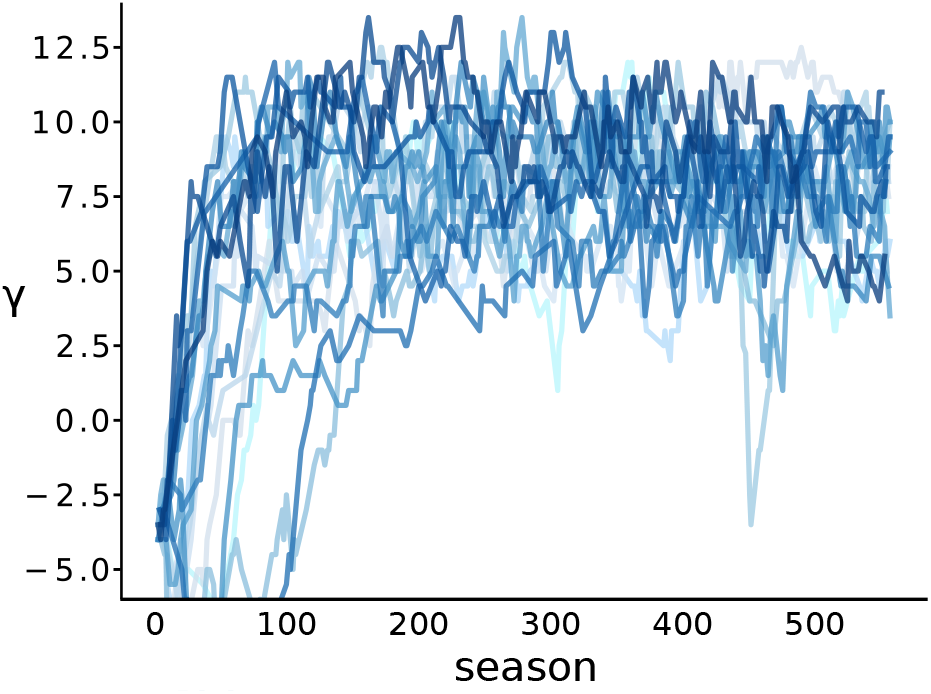
Evolution of adhesion in simulations starting from indivual evolved without adhesion. Evolution of adhesion in 20 simulations, started with 4 individuals that had evolved without adhesion (5 independent simulations per individual)

**Figure S5:**
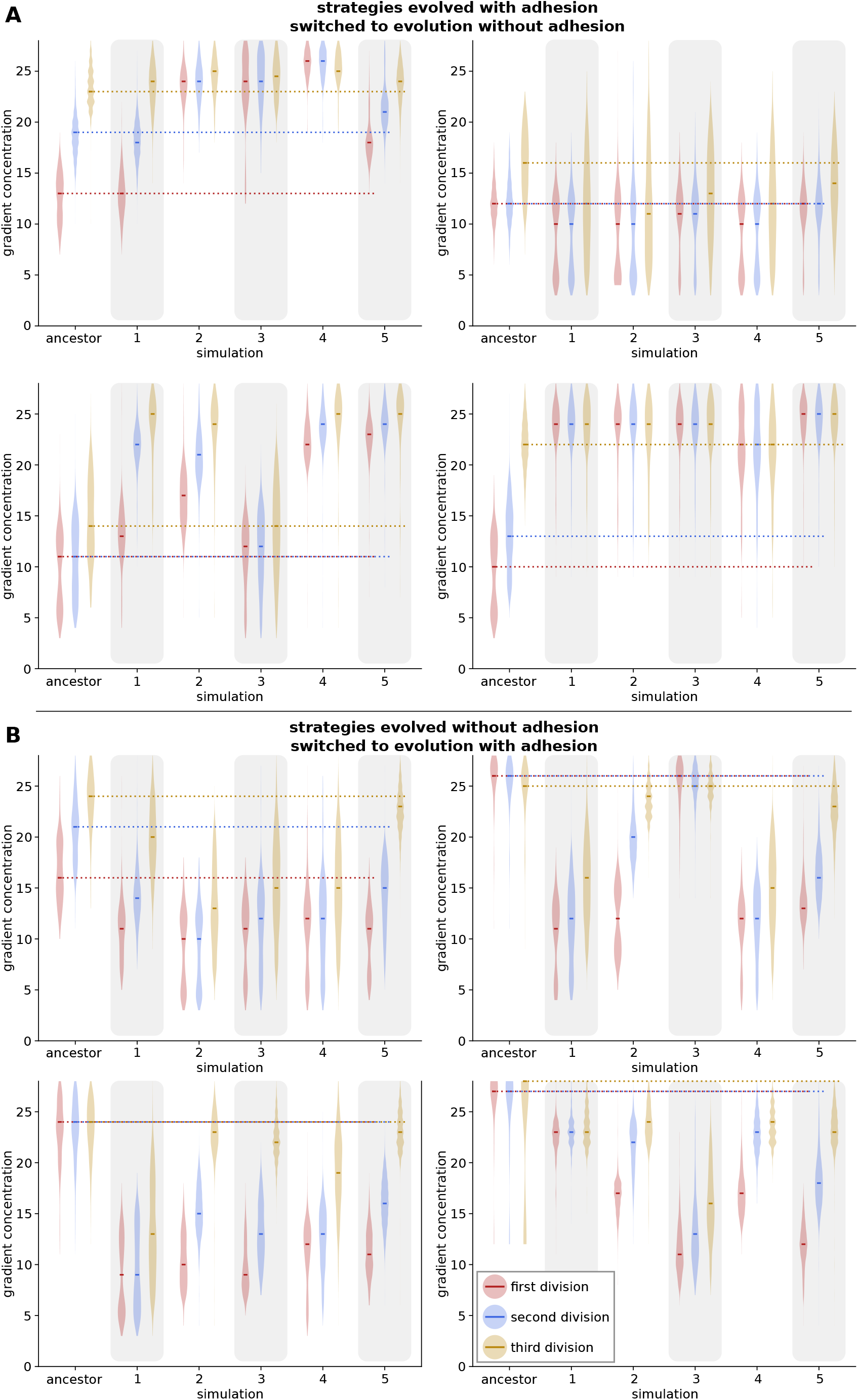
Evolution of gradient sensing in populations moved to opposite adhesion regime. Shown are the population distributions of the gradient level at which a division happens, in the ancestor and the five replicate simulations evolved from that ancestor.

**Figure S6:**
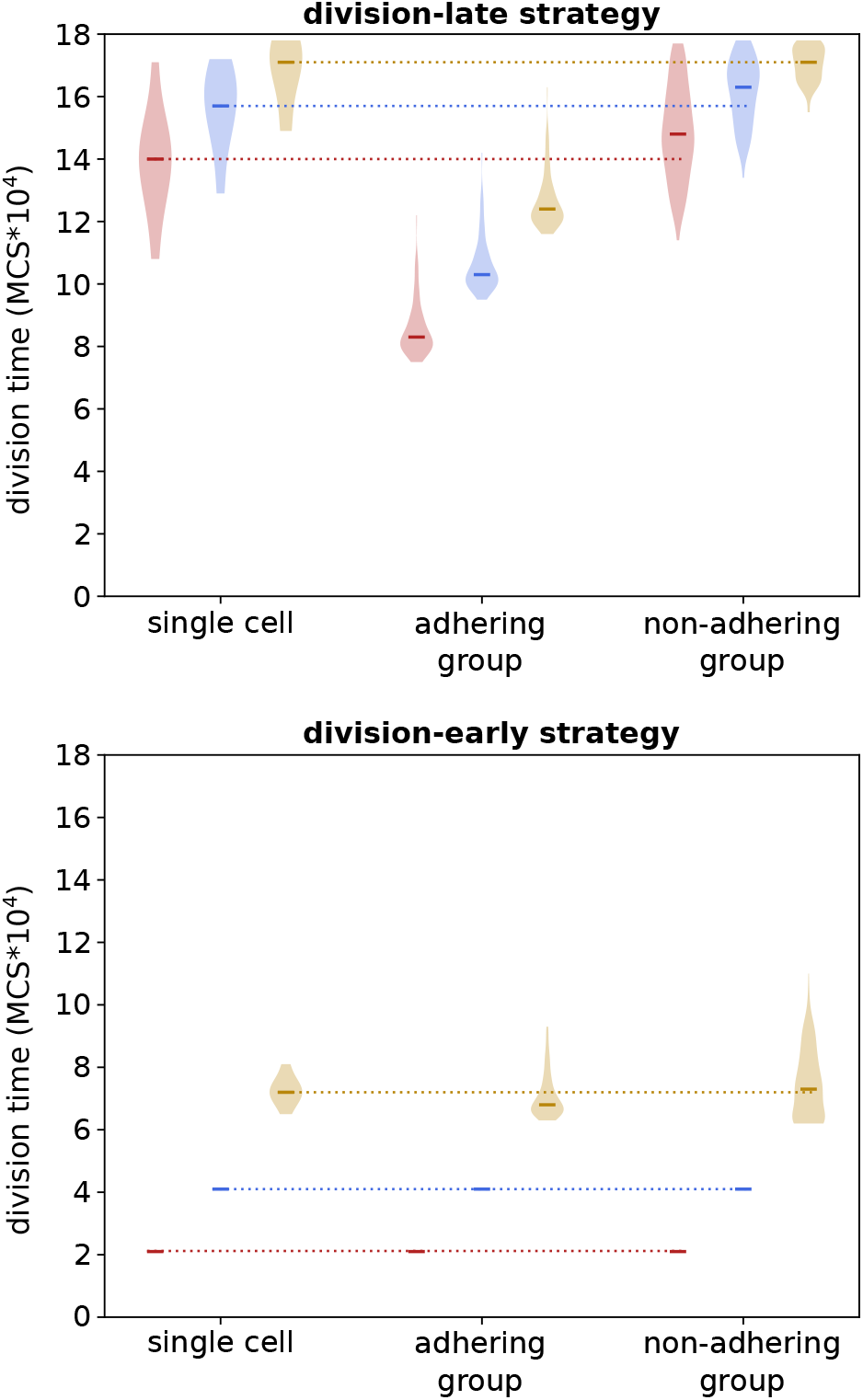
Effect of adhesion or non-adhesion on division timing. The distribution of division timings for simulations with a single cell compared to simulations of one season with 20 adhering or non-adhering cells. For simplicity, divisions were simulated by giving the cell the dividing state for the same amount of time, but no additional daughter cell was created. This kept the group size the same throughout the run. There were also no mutations of adhesion or regulation. We show here two example genomes, one evolved without adhesion, possessing a division-late strategy; and one evolved with adhesion, possessing a division-early strategy. Season duration=180000.

**Figure S7:**
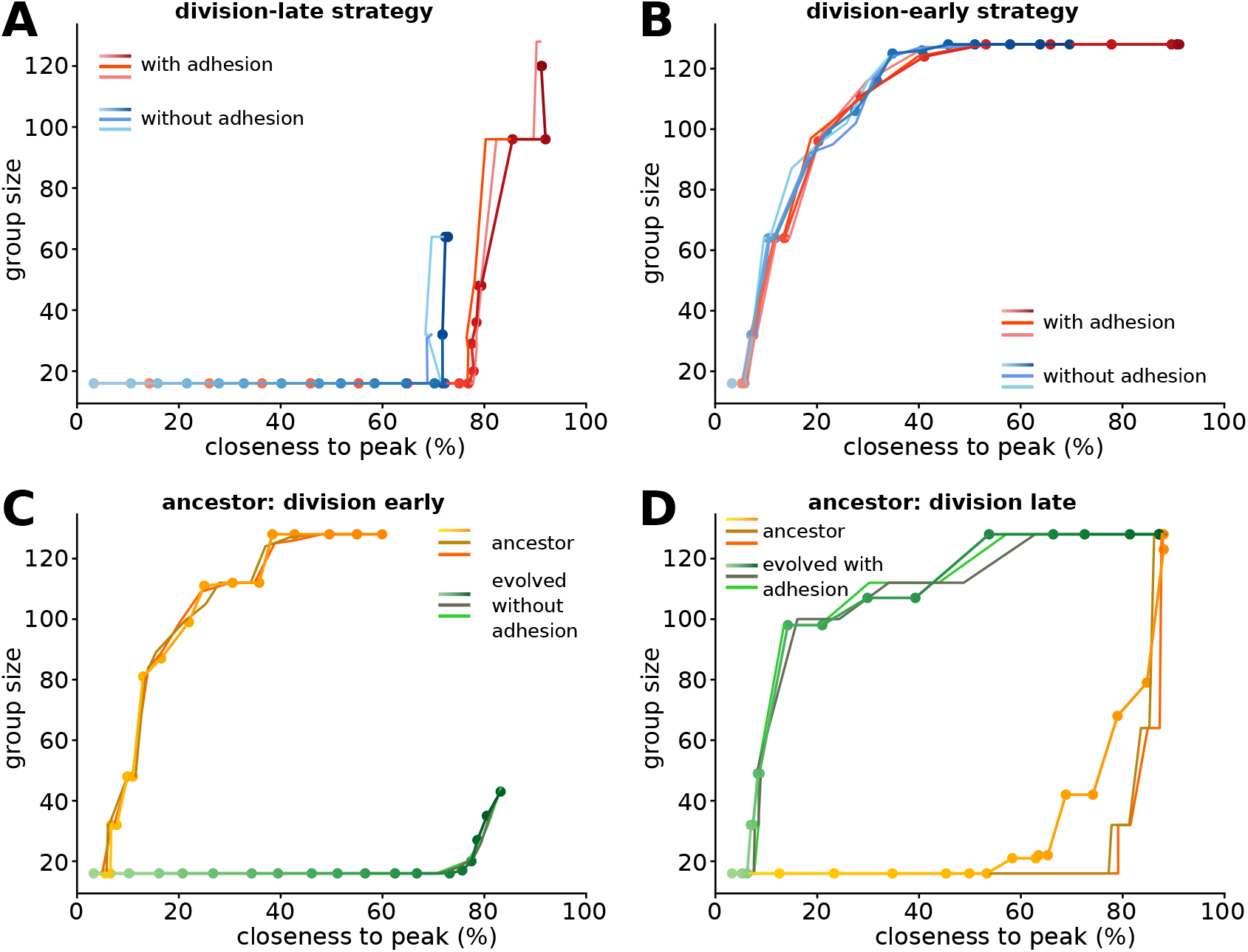
Replicates of competition experiment. Group size plotted against the median distance of cells to the peak of gradient, shown for the first season of the competition experiment (0%=maximum distance, 100%=at the peak). The graded lines with dots are the same as in figure 5. **a-b)** Competition between two groups with the same regulatory strategy, one adhering (*γ* = 6; a), the other non-adhering (*γ* = — 4; b). **c)** Competition between ancestral, division-early strategy and evolved, division-late strategy; both non-adhering. **d)** Competition between two adhering groups, one with an ancestral, division-late strategy and one with a strategy evolved after 500 seasons with possibility of adhesion (having become more division-early).

## Supplementary Videos

1. One season of a simulation which evolved a division-early strategy (Season duration=250000; high death rate; adhesion cannot evolve)

2. One season of a simulation which evolved a division-early strategy (Season duration=250000; high death rate; adhesion can evolve).

3. One season of a simulation which evolved a division-late strategy (Season duration=250000; high death rate; adhesion cannot evolve)

4. One season of a simulation which evolved a division-late strategy (Season duration=250000; high death rate; adhesion can evolve).

5. A competition experiment between a cluster of adhering cells and a cluster of non-adhering cells, with division-late strategy.

6. A competition experiment between a cluster of adhering cells and a cluster of non-adhering cells, with division-early strategy.

7. A competition experiment between a cluster of division-early ancestors and a cluster of division-late descendants – both nonadhering.

8. A competition experiment between a cluster of division-late ancestors and a cluster of division-early descendants – both adhering.

9. One season of a simulation with a clonal cluster of initially 50 cells (i.e. all have the same regulatory strategy). Individual regulatory strategy taken from the last season of a simulation in which the population was switched from non-adhering to evolution of adhesion.

## References

[1] John Maynard Smith and Eors Szathmary. The major transitions in evolution. Oxford, UK: WE Freeman, 1995.

[2] Richard K Grosberg and Richard R Strathmann. The evolution of multicellularity: a minor major transition? Annu. Rev. Ecol. Evol. Syst., 38:621–654, 2007.

[3] Thibaut Brunet and Nicole King. The origin of animal multicellularity and cell differentiation. Developmental cell, 43(2):124–140, 2017.

[4] Andrew H Knoll. The multiple origins of complex multicellularity. Annual Review of Earth and Planetary Sciences, 39:217–239, 2011.

[5] Martin E Boraas, Dianne B Seale, and Joseph E Boxhorn. Phagotrophy by a flagellate selects for colonial prey: a possible origin of multicellularity. Evolutionary Ecology, 12 (2):153–164, 1998.

[6] William C Ratcliff, R Ford Denison, Mark Borrello, and Michael Travisano. Experimental evolution of multicellularity. Proceedings of the National Academy of Sciences, 109(5): 1595–1600, 2012.

[7] Nicole King. The unicellular ancestry of animal development. Developmental Cell, 7(3): 313 – 325, 2004. ISSN 1534-5807.

[8] Stephen R. Fairclough, Zehua Chen, Eric Kramer, Qiandong Zeng, Sarah Young, Hugh M. Robertson, Emina Begovic, Daniel J. Richter, Carsten Russ, M. Jody Westbrook, Gerard Manning, B. Franz Lang, Brian Haas, Chad Nusbaum, and Nicole King. Premetazoan genome evolution and the regulation of cell differentiation in the choanoflagellate Salpingoeca rosetta. Genome Biology, 14(2):R15, February 2013. ISSN 1474-760X. doi: 10.1186/gb-2013-14-2-r15.

[9] Qingyou Du, Yoshinori Kawabe, Christina Schilde, Zhi-hui Chen, and Pauline Schaap. The evolution of aggregative multicellularity and cell–cell communication in the dictyostelia. Journal of molecular biology, 427(23):3722–3733, 2015.

[10] Núria Ros-Rocher, Alberto Pérez-Posada, Michelle M Leger, and Iñaki Ruiz-Trillo. The origin of animals: an ancestral reconstruction of the unicellular-to-multicellular transition. Open Biology, 11(2):200359, 2021.

[11] David L Kirk. A twelve-step program for evolving multicellularity and a division of labor. BioEssays, 27(3):299–310, 2005.

[12] Scott D Evans, Mary L Droser, and Douglas H Erwin. Developmental processes in ediacara macrofossils. Proceedings of the Royal Society B, 288(1945):20203055, 2021.

[13] Iaroslav Ispolatov, Martin Ackermann, and Michael Doebeli. Division of labour and the evolution of multicellularity. Proceedings of the Royal Society B: Biological Sciences, 279 (1734):1768–1776, 2011.

[14] Cristian A. Solari, John O. Kessler, and Raymond E. Goldstein. A general allometric and life-history model for cellular differentiation in the transition to multicellularity. The American Naturalist, 181(3):369–380, 2013. doi: 10.1086/669151. PMID: 23448886.

[15] Denis Tverskoi, Vladimir Makarenkov, and Fuad Aleskerov. Modeling functional specialization of a cell colony under different fecundity and viability rates and resource constraint. PLOS ONE, 13(8):1–27, 08 2018. doi: 10.1371/journal.pone.0201446.

[16] Andre Amado, Carlos Batista, and Paulo R.A. Campos. A mechanistic model for the evolution of multicellularity. Physica A: Statistical Mechanics and its Applications, 492: 1543–1554, 2018. ISSN 0378-4371. doi: https://doi.org/10.1016/j.physa.2017.11.080.

[17] Chikara Furusawa and Kunihiko Kaneko. Origin of multicellular organisms as an inevitable consequence of dynamical systems. The Anatomical Record, 268(3):327–342, 2002.

[18] Paulien Hogeweg. Evolving mechanisms of morphogenesis: on the interplay between differential adhesion and cell differentiation. Journal of Theoretical Biology, 203(4):317 – 333, 2000. ISSN 0022-5193.

[19] Leonardo Miele and Silvia De Monte. Aggregative cycles evolve as a solution to conflicts in social investment. PLoS computational biology, 17(1):e1008617, 2021.

[20] Merlijn Staps, Jordi van Gestel, and Corina E Tarnita. Emergence of diverse life cycles and life histories at the origin of multicellularity. Nature ecology & evolution, 3(8):1197–1205, 2019.

[21] Salva Duran-Nebreda, Adriano Bonforti, Raúl Montanez, Sergi Valverde, and Ricard Solé. Emergence of proto-organisms from bistable stochastic differentiation and adhesion. Journal of The Royal Society Interface, 13(117):20160108, 2016.

[22] ES Colizzi, B van Dijk, RMH Merks, DE Rozen, and RMA Vroomans. Evolution of genome fragility enables microbial division of labor. bioRxiv, 2021.

[23] Enrico Sandro Colizzi, Renske MA Vroomans, and Roeland MH Merks. Evolution of multicellularity by collective integration of spatial information. eLife, 9, 2020. doi: 10.7554/eLife.56349.

[24] Françaois Graner and James A. Glazier. Simulation of biological cell sorting using a two-dimensional extended potts model. Phys. Rev. Lett., 69:2013–2016, Sep 1992.

[25] James A. Glazier and François Graner. Simulation of the differential adhesion driven rearrangement of biological cells. Phys. Rev. E, 47:2128–2154, Mar 1993. doi: 10.1103/Phys-RevE.47.2128.

[26] Josephine T. Daub and Roeland M. H. Merks. Cell-Based Computational Modeling of Vascular Morphogenesis Using Tissue Simulation Toolkit, pages 67–127. Springer New York, New York, NY, 2015. ISBN 978-1-4939-1462-3.

[27] Jos Käfer, Takashi Hayashi, Athanasius FM Marée, Richard W Carthew, and François Graner. Cell adhesion and cortex contractility determine cell patterning in the drosophila retina. Proceedings of the National Academy of Sciences, 104(47):18549–18554, 2007.

[28] Susan D Hester, Julio M Belmonte, J Scott Gens, Sherry G Clendenon, and James A Glazier. A multi-cell, multi-scale model of vertebrate segmentation and somite formation. PLoS computational biology, 7(10):e1002155, 2011.

[29] Renske MA Vroomans, Paulien Hogeweg, and Kirsten HWJ ten Tusscher. Segments-pecific adhesion as a driver of convergent extension. PLoS computational biology, 11 (2):e1004092, 2015.

[30] Jordi van Gestel and Franz J. Weissing. Regulatory mechanisms link phenotypic plasticity to evolvability. Scientific Reports, 6(1):24524, Apr 2016. ISSN 2045-2322. doi: 10.1038/srep24524.

[31] Mark Zajac, Gerald L Jones, and James A Glazier. Simulating convergent extension by way of anisotropic differential adhesion. Journal of Theoretical Biology, 222(2):247–259, 2003.

[32] Antonis Rokas. The molecular origins of multicellular transitions. Current opinion in genetics & development, 18(6):472–478, 2008.

[33] Simon E Prochnik, James Umen, Aurora M Nedelcu, Armin Hallmann, Stephen M Miller, Ichiro Nishii, Patrick Ferris, Alan Kuo, Therese Mitros, Lillian K Fritz-Laylin, et al. Genomic analysis of organismal complexity in the multicellular green alga volvox carteri. Science, 329(5988):223–226, 2010.

[34] Daniel J Richter, Parinaz Fozouni, Michael B Eisen, and Nicole King. Gene family innovation, conservation and loss on the animal stem lineage. Elife, 7:e34226, 2018.

[35] Douglas H. Erwin. The origin of animal body plans: a view from fossil evidence and the regulatory genome. Development, 147(4), February 2020. ISSN 0950-1991, 1477-9129. doi: 10.1242/dev.182899.

[36] Gavriel Matt and James Umen. Volvox: A simple algal model for embryogenesis, morphogenesis and cellular differentiation. Developmental Biology, 419(1):99–113, November 2016. doi: 10.1016/j.ydbio.2016.07.014.

[37] Pauline Schaap. Evolutionary crossroads in developmental biology: Dictyostelium discoideum. Development, 138(3):387–396, 2011.

[38] Lana Strmecki, David M. Greene, and Catherine J. Pears. Developmental decisions in dictyostelium discoideum. Developmental Biology, 284(1):25–36, 2005. ISSN 0012-1606. doi: https://doi.org/10.1016/j.ydbio.2005.05.011.

[39] Arielle Woznica, Alexandra M Cantley, Christine Beemelmanns, Elizaveta Freinkman, Jon Clardy, and Nicole King. Bacterial lipids activate, synergize, and inhibit a developmental switch in choanoflagellates. Proceedings of the National Academy of Sciences, 113 (28):7894–7899, 2016.

[40] Arielle Woznica, Joseph P. Gerdt, Ryan E. Hulett, Jon Clardy, and Nicole King. Mating in the closest living relatives of animals is induced by a bacterial chondroitinase. Cell, 170(6):1175–1183.e11, 2017. ISSN 0092-8674. doi: https://doi.org/10.1016/j.cell.2017.08.005.

[41] José Muñoz-Dorado, Francisco J Marcos-Torres, Elena García-Bravo, Aurelio Moraleda-Muñoz, and Juana Pérez. Myxobacteria: moving, killing, feeding, and surviving together. Frontiers in microbiology, 7:781, 2016.

[42] Christopher Toret, Andrea Picco, Micaela Boiero-Sanders, Alphee Michelot, and Marko Kaksonen. The cellular slime mold fonticula alba forms a dynamic, multicellular collective while feeding on bacteria. Current Biology, 2022.

[43] Thomas Garcia, Leonardo Gregory Brunnet, and Silvia De Monte. Differential adhesion between moving particles as a mechanism for the evolution of social groups. PLoS computational biology, 10(2):e1003482, 2014.

[44] Jaideep Joshi, Iain D Couzin, Simon A Levin, and Vishwesha Guttal. Mobility can promote the evolution of cooperation via emergent self-assortment dynamics. PLoS computational biology, 13(9):e1005732, 2017.

[45] Kirill V. Mikhailov, Anastasiya V. Konstantinova, Mikhail A. Nikitin, Peter V. Troshin, Leonid Yu. Rusin, Vassily A. Lyubetsky, Yuri V. Panchin, Alexander P. Mylnikov, Leonid L. Moroz, Sudhir Kumar, and Vladimir V. Aleoshin. The origin of metazoa: a transition from temporal to spatial cell differentiation. BioEssays, 31(7):758–768, 2009. doi: https://doi.org/10.1002/bies.200800214.

[46] Renske MA Vroomans and Enrico Sandro Colizzi. cellevol 2.0: Tsto-based software and scripts for studying the evolution of differentiated multicellularity. 2022. URL https://github.com/RenskeVroomans/regulatiom_evolution.

[47] James A Glazier, Ariel Balter, and Nikodem J Popławski. Magnetization to morphogenesis: a brief history of the glazier-graner-hogeweg model. In Single-Cell-Based Models in Biology and Medicine, pages 79–106. Springer, 2007.

[48] Leonie van Steijn, Inge M. N. Wortel, Clément Sire, Loïc Dupré, Guy Theraulaz, and Roeland M. H. Merks. Computational modelling of cell motility modes emerging from cell-matrix adhesion dynamics. PLOS Computational Biology, 18(2):1–28, 02 2022. doi: 10.1371/journal.pcbi.1009156. URL https://doi.org/10.1371/journal.pcbi.1009156.

[49] Joost B. Beltman, Athanasius F.M. Marée, Jennifer N. Lynch, Mark J. Miller, and Rob J. de Boer. Lymph node topology dictates T cell migration behavior. The Journal of Experimental Medicine, 204(4):771–780, 03 2007. ISSN 0022-1007.

